# Systems vaccinology analysis of a recombinant vaccinia-based vector reveals diverse innate immune signatures at the injection site

**DOI:** 10.1101/2020.08.17.254938

**Authors:** Jessamine E. Hazlewood, Troy Dumenil, Thuy T. Le, Andrii Slonchak, Stephen H. Kazakoff, Ann-Marie Patch, Lesley-Ann Gray, Paul M. Howley, Liang Liu, John D. Hayball, Kexin Yan, Daniel J. Rawle, Natalie A. Prow, Andreas Suhrbier

## Abstract

Poxvirus systems have been extensively used as vaccine vectors. Herein a systems vaccinology analysis of intramuscular injection sites provides detailed insights into host innate immune responses, as well as expression of vector and recombinant immunogen genes, after vaccination with a new multiplication defective, vaccinia-based vector, Sementis Copenhagen Vector. Chikungunya and Zika virus immunogen mRNA and protein expression was associated with necrosing skeletal muscle cells surrounded by mixed cellular infiltrates. Adjuvant signatures at 12 hours post-vaccination were dominated by TLR3, 4 and 9, STING, MAVS, PKR and the inflammasome. Th1 cytokine signatures were dominated by IFNγ, TNF and IL1β, and chemokine signatures by CCL5 and CXCL12. Multiple signatures associated with dendritic cell stimulation were evident. By day seven, vaccine transcripts were absent, and cell death, neutrophil, macrophage and inflammation annotations had abated. No compelling arthritis signatures were identified. Such innate systems vaccinology approaches should inform refinements in poxvirus-based vector design.

## Introduction

A range of vaccine vector systems based on vaccinia virus (VACV) and other poxviruses have been developed, with several sold as products and many more in development and in human clinical trials (Prow *et al*., 2018a). These include Modified Vaccinia Ankara (MVA) (Koch *et al*., 2020; Sutter, 2020), NYVAC (Pantaleo *et al*., 2019), ALVAC (Laher *et al*., 2020), fowlpox (Gatti-Mays *et al*., 2019), LC16m8 (Omura *et al*., 2018), and raccoonpox (Stading *et al*., 2017). A large series of recombinant MVA (rMVA) vaccines have been evaluated in non-human primate (NHP) studies (Nagata *et al*., 2018) and in human clinical trials (Pittman *et al*., 2019; Prow *et al*., 2018a), with MVA licensed as a smallpox vaccine (sold as Imvanex/Imvamune). Recombinant poxvirus vector systems have a number of attractive features for vaccine development including a large payload capacity (at least 25,000 base pairs), potential for cold chain-independent distribution, lack of vaccine DNA integration and induction of both cellular and humoral immunity (Prow *et al*., 2018a). Nevertheless, a range of strategies are being sought to improve immunogenicity and reduce reactogenicity (Albarnaz *et al*., 2018; Chea *et al*., 2019; Izzi *et al*., 2014; Joachim *et al*., 2020; Koch *et al*., 2020; Marin *et al*., 2018). Both these key characteristics of vaccines are largely dictated by the early behavior of the vaccine at the injection site. However, a comprehensive systems vaccinology approach to characterize the post-inoculation injection site responses has not been undertaken for a recombinant poxvirus vaccine.

The Sementis Copenhagen Vector (SCV), derived from the Copenhagen strain of VACV, was recently described (Eldi *et al*., 2017; Prow *et al*., 2018b). SCV can replicate its DNA but is rendered unable to generate viral progeny in vaccine recipients by virtue of a targeted deletion of the *D13L* gene that encodes the essential viral assembly protein, D13. Recombinant SCV vaccines are produced in Chinese Hamster Ovary (CHO) cells modified to express D13 and the host range protein, CP77 (Eldi *et al*., 2017). A single construct recombinant SCV vaccine encoding the structural gene cassettes of both chikungunya virus (CHIKV) and Zika virus (ZIKV) (SCV-ZIKA/CHIK) was constructed with each polyprotein immunogen driven by the same synthetic strong early late promoter (Alharbi, 2019), but from two distinct distant loci from within the SCV genome (Prow *et al*., 2018b). A dual ZIKV and CHIKV vaccine was deemed attractive as these virus co-circulate in overlapping geographic regions, and can co-infect both mosquitoes and humans (Prow *et al*., 2020; Schrauf *et al*., 2020; Suhrbier, 2019). SCV-ZIKA/CHIK was shown to protect against CHIKV and ZIKV in a series of mouse models (Prow *et al*., 2018b). In NHPs the vaccine also induced neutralizing antibodies against VACV, CHIKV and ZIKV and provided protection against ZIKV viremia (Prow *et al*., 2020).

Systems vaccinology uses mRNA expression profiling to gain a detailed molecular understanding of the behavior of vaccines *in vivo*, thereby informing design and development (Sharma *et al*., 2019). The approach has been used to understand and predict immunogenicity (Matthijs *et al*., 2019; Natrajan *et al*., 2019), reactogenicity/safety (Gonzalez-Dias *et al*., 2020; McKay *et al*., 2019) and adjuvant activity (Ng *et al*., 2019b; Sarkar *et al*., 2019). Most systems vaccinology studies have analyzed peripheral blood post vaccination, as this is readily accessible in humans. However, herein we described RNA-Seq and bioinformatics analyses of injection sites after intramuscular (i.m.) vaccination. Adult wild-type were vaccinated with SV-ZIKA/CHIK and muscles were harvested at 12 hours post vaccination to characterize early injections site innate responses and identify adjuvant signatures. As vector-induced cytopathic effects (CPE) are only just beginning (at least *in vitro*) at 12 hours post infection (Eldi *et al*., 2017), this was also deemed a suitable time to investigate expression *in vivo* of both viral vector genes and expression of the recombinant immunogens. Muscle tissue was similarly analysed on day 7 post i.m. vaccination to determine the persistence of vaccine transcripts and characterize the evolution of injection site inflammatory responses at a time when vaccine-induced adaptive immune responses are being generated. This is also the time when inflammatory lesions develop after VACV vaccination (Darling *et al*., 2014; Frey *et al*., 2002; Fulginiti *et al*., 2003; He *et al*., 2014; Parrino *et al*., 2007; Talbot *et al*., 2006; Tian *et al*., 2017). Finally, feet were harvested on day 7 post vaccination to determine whether SCV-ZIKA/CHIK vaccination might be associated with an arthritic signature, given that a previous live-attenuated CHIKV vaccine induced arthralgia (Edelman *et al*., 2000). The characterization provided herein of vaccine gene expression and innate host responses at the injection site provide both a process and insights that may inform future endeavors to improve immunogenicity whilst limiting reactogenicity of poxvirus-based vaccine vectors.

## Results

### RNA-Seq and differential gene expression

Mice were vaccinated i.m with SCV-ZIKA/CHIK or were mock vaccinated with PBS; feet and quadriceps muscles were then harvested at 12 hours and 7 days post vaccination (S1A Fig). Each of the 3 biological replicates comprised pooled RNA from 4 feet or 4 quadriceps muscles from 4 different mice (S1B Fig). Poly-adenylated mRNA was sequenced using the Illumina HiSeq 2500 Sequencer. Per base sequence quality for >93% bases was above Q30 for all samples. The mean total paired-end reads per group ranged from ≈19 to 24 million, with >91.6% of reads mapping to the mouse genome (S1C Fig). Five groups were analyzed in triplicate (i) quadriceps muscles from mock vaccinated mice (MQ), (ii) quadriceps muscles from mice vaccinated with SCV-ZIKA/CHIK taken 12 hours post vaccination (SCV12hQ), (iii) quadriceps muscles from mice vaccinated with SCV-ZIKA/CHIK and taken 7 days post vaccination (SCVd7Q), (iv) feet taken from mice mock vaccinated i.m. (MF) and (iv) feet from SCV-ZIKA/CHIK vaccinated mice taken 7 days post vaccination (SCVd7F). Reads were mapped to the *Mus musculus* genome (mm10) using STAR aligner, with a similar distribution of read counts observed for all samples (S1D Fig). Differentially expressed genes were generated for MQ vs SCV12hQ (for early post-vaccination injection site responses), MQ vs SCVd7Q (for injection site responses on day 7 post-vaccination) and MF vs SCVd7F (to evaluate potential arthritogenic side effects associated with vaccination) (Smear plots are provided in S1E Fig).

### Read alignments to the SCV-ZIKA/CHIK vaccine genome

Expression of the vaccine vector and recombinant ZIKA and CHIK immunogen genes was analysed by aligning reads to a combined reference that included mouse, VACV, ZIKV and CHIKV genomes. Given vaccine transcripts can only be expressed in host cells, SCV-ZIKA/CHIK vaccine reads were expressed as a percentage of total RNA sequencing reads mapping to the mouse genome (Fig. 1A; S1A Table). The overall expression profile remained similar when an alternative aligner was used (S2 Fig). The only time at which significant vaccine-derived reads were evident was in quadriceps muscles at 12 hours post vaccination (Fig. 1A; S1A,B Table), suggesting that the vaccine had largely been cleared from the injection site by day 7. This is consistent with *in vitro* data showing that SCV induces cytopathic effects (CPE) in infected cells within a few days and that SCV is unable to produce viral progeny (Eldi *et al*., 2017; Prow *et al*., 2018a; Prow *et al*., 2018b). The paucity of vaccine reads in the feet 7 days post vaccination (Fig. 1A, SCVd7Q) also illustrates that the vaccine does not disseminate to and/or persist in joint tissues (a potential safety concern; see below). The percentage of reads mapping to a murine house-keeping gene, RPL13A (Schroder *et al*., 2010), was similar for the 3 samples from quadriceps muscles, and for the two samples from feet (Fig. 1B), illustrating that the low vaccine read counts for mock and day 7 samples (Fig. 1A) were not due to low read counts generally for those samples.

**Figure 1.**
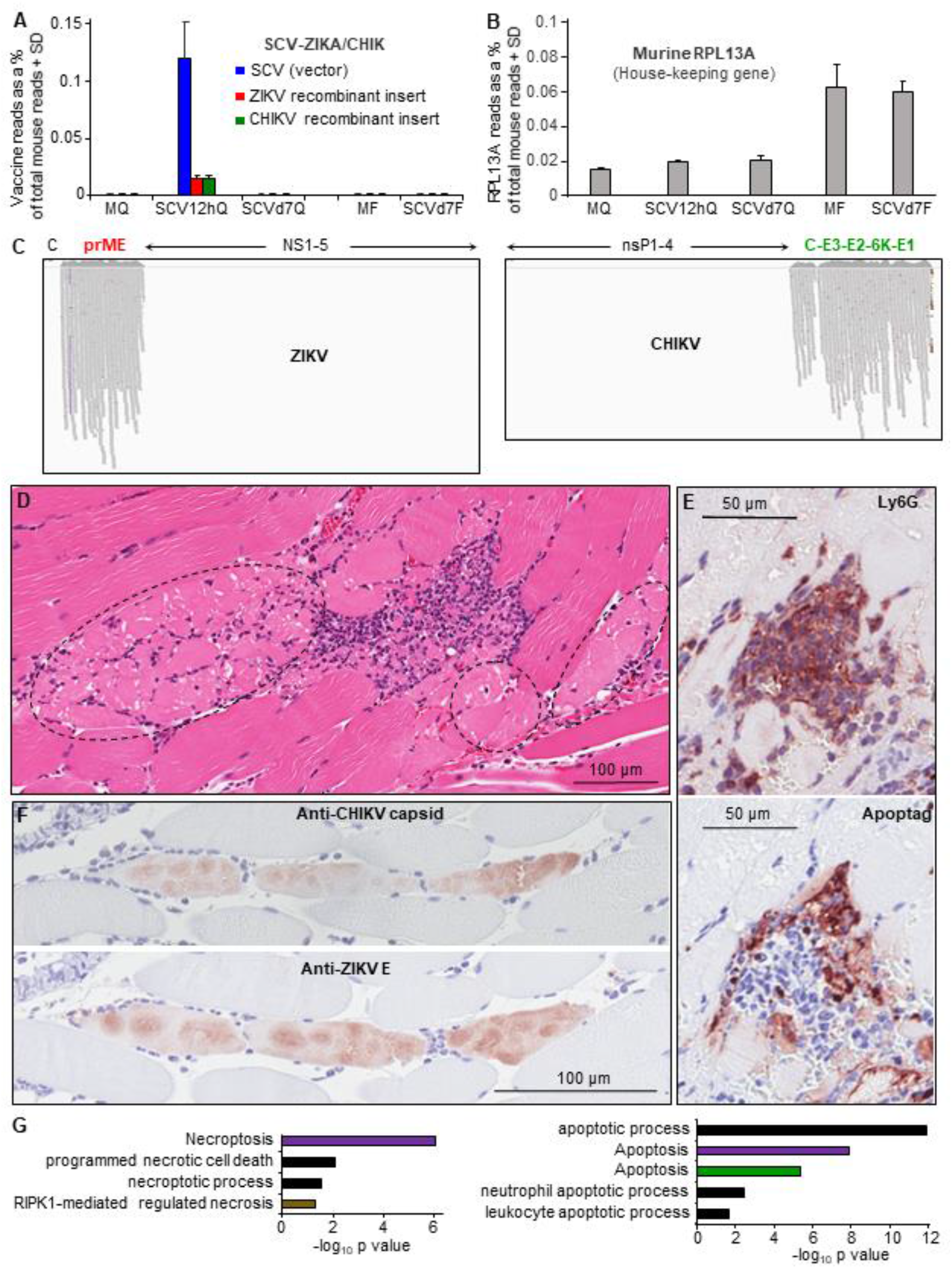
Read alignments to the viral genomes. (A) RNA-Seq reads from each of the five groups aligned to the three viral genomes (the vector, SCV, and the two recombinant immunogen inserts from ZIKV and CHIKV); MQ - quadriceps muscles from mock vaccinated, SCV12hQ mice quadriceps muscles from SCV-ZIKA/CHIK vaccinated mice 12 hours post vaccination, SCVd7Q -quadriceps muscles from SCV-ZIKA/CHIK vaccinated mice taken 7 days post vaccination, MF – feet from mock vaccinated mice 7 days post vaccination, and SCVd7F - feet from SCV-ZIKA/CHIK vaccinated mice 7 days post vaccination. The number of viral reads is expressed as a percentage of the number of reads mapping to the mouse genome, with 3 biological replicates providing the SD (Supplementary Fig. 1). The bars plotting to ≈0% had values ranging from 0 to 3.5×10^−5^ %. (B) RNA-Seq reads from each of the five groups aligned to the house-keeping gene, RPL13A, also expressed as a percentage of the number of reads mapping to the mouse genome. (C) IGV visualization of reads aligned to the recombinant structural polyprotein immunogens of ZIKV (prME) and CHIKV (C-E3-E2-6K-E1), which are encoded in the SCV-ZIKA/CHIK vaccine. All reads from all replicates are shown (for details see (S1A Table). As expected, no reads mapped to the non-structural genes of ZIKV or CHIKV (NS1-5 and nsP1-4, respectively), as these are not encoded in SCV-ZIKA/CHIK. (Vertical purple lines for ZIKV indicate base call errors after a string of Gs). (Reads mapping to the SCV genome are shown in S1B Table). (D) H&E staining of injection site 12 hours post vaccination. Dotted ovals indicate muscle cells in early stages of necrosis (pink staining). (E) Top; IHC with anti-Ly6G staining for neutrophils (parallel section to D focusing on area of infiltrates). Bottom: Apoptag staining of the same area, illustrating apoptosis within areas of infiltrating cells. (F) Top: IHC for CHIK capsid protein. Bottom: parallel section showing IHC for ZIKA E protein. (G) Cell death annotation from Cytoscape analysis of up-regulated DEGs at 12 hours post vaccination (MQ vs SCV12Q) (S2C Table) divided into non-apoptotic signatures (left) and apoptotic signatures (right). KEGG Pathways (purple), Go process (black), Reactome Pathways (brown), UniProt Keywords (green).

A criticism of virally vectored vaccines has been that viral vector transcripts can be so much more abundant than recombinant immunogen transcripts, resulting in vaccine responses excessively biased towards vector proteins (Harrington *et al*., 2002; Tscharke *et al*., 2005; Wyatt *et al*., 2017). However, ≈20% of all the SCV-ZIKA/CHIK vaccine reads mapped to the two recombinant immunogens, even though the ZIKV and CHIKV structural protein sequences in the vaccine genome were relative small (2067 bp and 3747 bp, respectively), when compared to the large SCV genome (≈190,000 bp). This perhaps attests to the strength of the poxvirus synthetic strong early late promoter (Alharbi, 2019) used for CHIKV and ZIKV immunogens in the SCV-ZIKA/CHIK vaccine (Prow *et al*., 2018b).

Expression of two immunogens in a single poxvirus vector construct carries the risk that one immunogen is expressed significantly better than the other, a problem encountered in a variety of settings (Prow *et al*., 2018a). A comparable number of reads mapped to the recombinant CHIK and ZIKA inserts (Fig. 1A), with these two inserts distantly separate from each other in the SCV genome and driven from the same promoter (Prow *et al*., 2018a). This approach would seem largely to ensure (at least in SCV-ZIKA/CHIK) that comparable levels of mRNA are produced for each of the two immunogens.

Reads aligned to the CHIKV and ZIKV genomes were viewed using Integrative Genome Viewer (IGV) (Robinson *et al*., 2011). As expected, reads mapped to prME and C-E3-E2-6K-E1, which are encoded by SCV-ZIKA/CHIK; but not ZIKV capsid nor the non-structural proteins from both arboviruses (NS1-5 and nsP1-4), which are not encoded by SCV-ZIKA/CHIK (Fig. 1C). Premature immunogen termination has been described previously for a VACV-based vaccine (Earl *et al*., 1990), with VACV transcription occurring in the cytoplasm (Moss *et al*., 1991). No evidence for premature termination of SCV-ZIKA/CHIK immunogen transcription was apparent (Fig. 1C).

Read alignments to genes encoded by SCV are described and annotated in detail in S1B Table, with immune-modulation and cell-death modulating proteins highlighted, along with annotations regarding their activity in mice and their activity in the Copenhagen strain of VACV, from which SCV was derived. Many of these genes are referred to below.

### Injection site histology and immunohistochemistry at 12 hours post vaccination

H&E staining of the intramuscular injection sites showed that some skeletal muscle cells displayed fragmented pale cytoplasm with loss of striation and small condensed pyknotic nuclei indicative of necrosis (Fig. 1D, dotted ovals). These necrotic cells were partially surrounded by mixed inflammatory cells infiltrates (high densities of purple nuclei) and some cellular debris (Fig. 1D.

Immunohistochemistry (IHC) with a neutrophil-specific marker, anti-Ly6G (Poo *et al*., 2014; Prow *et al*., 2019), illustrated that the infiltrates contained abundant neutrophils (Fig. 1E, top panel, Ly6G). Interestingly, neutrophils have recently been shown to contribute to adjuvant activity (Stephen *et al*., 2017). The infiltrates also contained areas staining with Apoptag indicating apoptosis (Fig. 1E, parallel section, bottom panel, Apoptag).

IHC with monoclonal antibodies recognizing CHIKV capsid (5.5G9 (Goh *et al*., 2015) and ZIKV envelope (4G2) (Hobson-Peters *et al*., 2019), clearly illustrated expression of vaccine antigens in skeletal muscle cells 12 hours post infection (Fig. 1F). The spherical/oval cytoplasmic staining patterns likely reflect the well described cytoplasmic factories wherein the poxvirus coordinates protein expression and subjugates host functions (Katsafanas *et al*., 2007; Kieser *et al*., 2020; Paszkowski *et al*., 2016). These results illustrated that the immunogen mRNA expression seen in Fig. 1A and C translates into protein expression in muscle cells *in vivo*.

### Injection site host cell death signatures at 12 hours post vaccination

The mode of cell death for a host cell expressing vaccine immunogens can have important implications for immunogenicity, with necrosis often favored over apoptosis (Chea *et al*., 2019; Gargett *et al*., 2014). RNA-Seq analysis of the mouse i.m. injection sites 12 hours post vaccination (MQ vs SCV12hQ; full gene list in S2A Table) provided a set of differentially expressed genes (DEGs) (S2B Table; FDR or q ≤0.01, fold change ≥2 and sum of all counts across the six samples >6). The up-regulated DEGs (n=1390; S2B Table) were analyzed by Cytoscape (S2C Table), with cell death terms suggesting the presence of apoptosis, necroptosis and necrosis (Fig. 1G). Skeletal muscle cells are generally resistant to apoptosis (Schwartz, 2008, 2018) and VACV’s apoptosis inhibitor, B13R (Chea *et al*., 2019), was also expressed at the injection site (S1B Table). Skeletal muscle cells have recently been shown to be able to undergo necroptosis (Morgan *et al*., 2018), with skeletal muscle necrosis well described (Lentscher *et al*., 2020; Szugye, 2020). The apoptosis signatures (Fig. 1G) may largely be associated with infiltrating leukocytes such as neutrophils, which are highly prone to apoptosis (Soehnlein *et al*., 2010), as Apoptag staining was clearly present in the aforementioned infiltrates (Fig. 1E, Apoptag). MVA can induce apoptosis *in vitro* and in certain settings *in vivo* (Chea *et al*., 2019; Torres-Domínguez *et al*., 2019) and SCV can induce apoptosis (at least *in vitro*, S3 Fig); however, the mode of cell death elucidated *in vitro* may not be recapitulated in primary skeletal muscle cells *in vivo*.

### Large Toll-like receptor signatures at 12 hours post vaccination

The up-regulated DEGs (for MQ vs SCV12hQ; S2B Table) analyzed as above by Cytoscape returned multiple terms associated with innate immune responses (S2C Table). To provide insights into the early innate host immune responses and potential adjuvant signatures induced by SCV-ZIKA/CHIK vaccination, the full DEG list (1608 genes) for MQ vs SCV12hQ (S2B Table) was analyzed by Ingenuity Pathway Analysis (IPA) using the Up-Stream Regulator (USR) function and the direct and indirect interaction option. The list of USRs (S2D Table) illustrated a highly significant Toll-like receptor (TLR) signature, dominated by TLR3 and 4, followed by TLR9, 7 and 2 (Fig. 2A). Although other TLRs (TLR1, 5, 6, 7/8, 8) were also identified, the number of unique DEGs responsible for these annotations was low (Fig. 2A, numbers in brackets), arguing these were less reliable USRs as they arose from subsets of DEGs already used in the annotations for TLR3, 4, 9, 7 and/or 2 (S2D Table, Target molecules in dataset). Given the common signaling pathways used by all TLRs, primarily involving MyD88 and/or TICAM1/TRIF, overlap in genes induced via the different TLRs would be expected.

**Figure 2.**
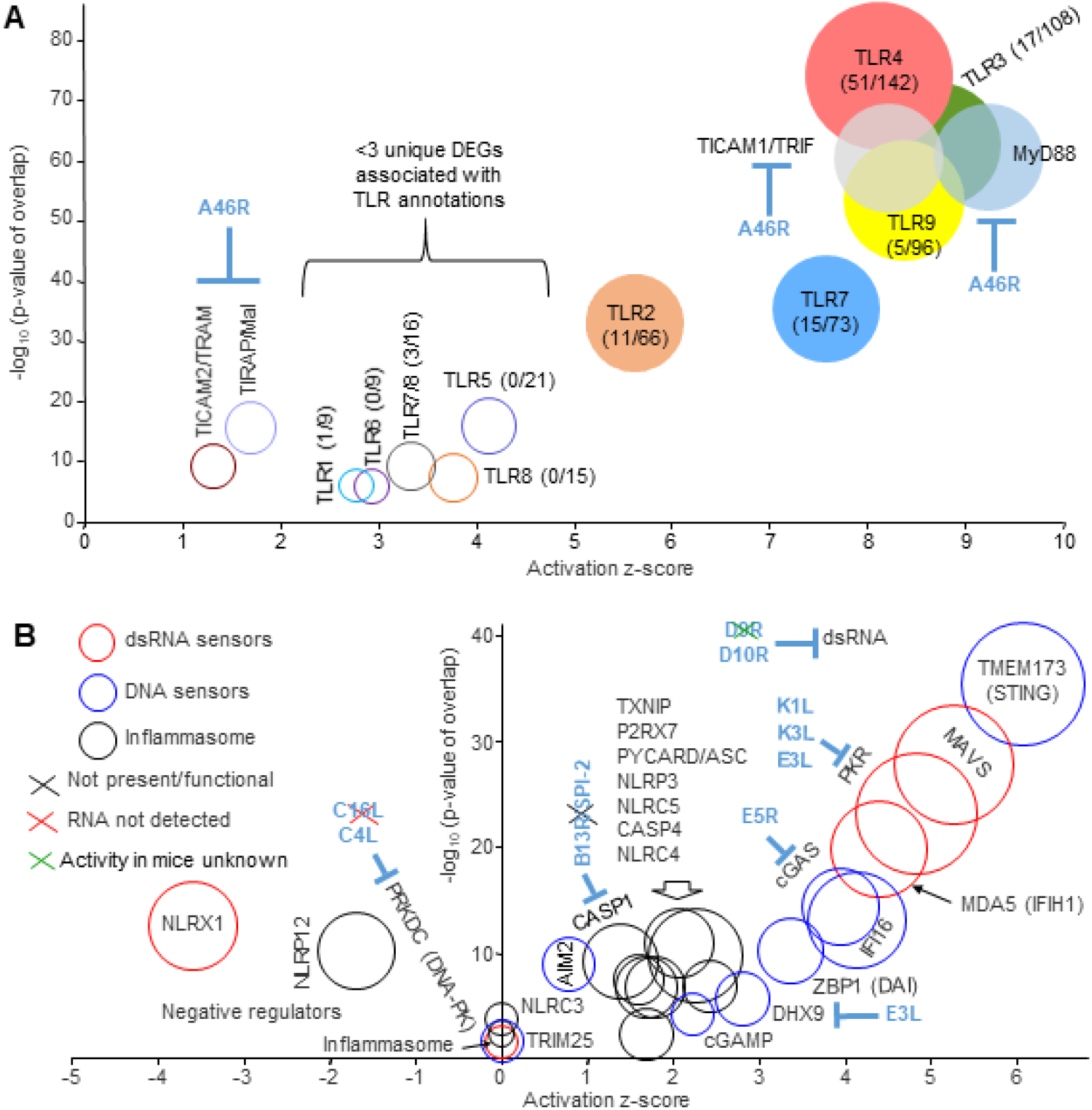
TLR and cytosolic sensor signatures at 12 hours post vaccination. (A) TLR signatures identified by IPA USR analysis (S2D Table) of 1608 DEGs identified in quadriceps muscles 12 hours post SCV-ZIKA/CHIK vaccination (MQ vs SCV12hQ; S2B Table)). Circle diameters reflect the number of DEGs associated with each IPA USR annotation. Numbers in brackets indicate the number of unique DEGs associated with each annotation over the total number of DEGs associated with the TLR annotation; circles with colored fills contain >3 DEGs uniquely associated with the indicated TLR annotation. A46R is expressed in the cytoplasm of infected cells. (B) Cytosolic sensor signatures identified by IPA USR analysis (S2D Table). Sensors divided into 3 categories associated with dsRNA (red circles), DNA (blue circles) and inflammasome activation (black circles). Circle diameters reflect the number of DEGs associated with each annotation. VACV genes encoding cytoplasmic inhibitors are shown in blue, with the black cross indicating that the gene/protein is not present or not functional in SCV (or in the Copenhagen strain of VACV), the red cross indicating that the gene was not detected by RNA-Seq of MQ vs SCV12hQ, the green cross indicate that the activity in mice is unknown (see S1B Table).

The identification herein of TLR4, TLR3, TLR9, TLR7 and TLR2 signatures (Fig. 2A) is notably consistent with the literature on poxvirus infections in GMO mice (summarized in S2E Table). TLR4 is stimulated by an unknown ligand present in/on VACV particles, with TLR4 required for effective antiviral activity and protection against mortality in mice after VACV infection (Hutchens *et al*., 2008b). TLR3 stimulation is likely mediated by dsRNA derived from the abundant complementary RNA transcripts produced late in the VACV infection cycle (Wolferstatter *et al*., 2014). TLR3 stimulation in VACV-infected mice promotes inflammatory cytokine production and immunopathology (Hutchens *et al*., 2008a). TLR7 (which detects ssRNA) is expressed on plasmacytoid dendritic cells and B cells, with TLR7 and TLR9 important for type I interferon secretion by dendritic cells following fowlpox infection (Lousberg *et al*., 2011). TLR9 is required for survival of mice following ectromelia virus infections (Samuelsson *et al*., 2008; Sutherland *et al*., 2011) and is likely stimulated by viral unmethylated ssDNA containing CpG motifs (Li *et al*., 2010) and/or mitochondrial DNA (Wang *et al*., 2017) released by viral CPE. TLR2 stimulation during VACV infection in mice has minimal impact on viral replication (Davies *et al*., 2014), but does promote NK activation and CD8 T cell expansion and memory (Martinez *et al*., 2010; Quigley *et al*., 2009). Thus both SCV and VACV would appear to stimulate TLR2, whereas MVA is reported not to do so (Price *et al*., 2014). To the best of our knowledge, there is no literature suggesting a role for TLR1, 5, 6 or 7 in poxvirus infections, consistent with the low number of unique DEGs for these annotations (Fig. 2A). The role of TLR8 in VACV infections remains controversial (Bauer *et al*., 2010), with TLR8 nonfunctional in mice (Ng *et al*., 2019a).

VACV produces an inhibitor of TRIF, MYD88, TRAM and MAL, called A46 or VIPER (encoded by A46R), a protein reported to be active in murine systems (Lysakova-Devine *et al*., 2010). However, A46 is expressed in the cytoplasm of SCV-infected cells and not in neighboring uninfected cells that may also express TLRs. Such cells might sense TLR agonists comprising viral pathogen-associated molecular patterns (PAMPs) and/or damage-associated molecular patterns (DAMPs) released by SCV infection-induced CPE (Eldi *et al*., 2017; Ink *et al*., 1995; Tsung *et al*., 1996).

### Multiple cytoplasmic sensor signatures at 12 hours post vaccination

The IPA analysis of DEGs for MQ vs SCV12hQ (S2D Table, direct and indirect) produced a series of USRs associated with (i) detection of cytoplasmic dsRNA (Fig. 2B, red circles) via MAVS, MDA5 and PKR, (ii) detection of cytoplasmic DNA (Fig. 2B, blue circles) dominated by STING/IFI16/cGAS, and (iii) activation of the inflammasome (Fig. 2B, black circles). These results again show marked consistency with existing poxvirus literature on KO mouse infections (summarized in S2E Table). dsRNA from complementary VACV RNA transcripts stimulates MAVS signaling (Myskiw *et al*., 2011), likely via MDA5 (Deng, 2017). Stimulation of MDA5 or RIG-I and PKR by VACV *in vitro* has been reported previously (Myskiw *et al*., 2011; Pichlmair *et al*., 2009), with both MAVS and MDA5 reported to contribute to host defense against VACV infection (Deng, 2017). PKR activation is also enhanced by MDA5 (Pham *et al*., 2016). Like SCV, the canarypox virus vector, ALVAC, also stimulates the cGAS/IFI16/STING pathway (Liu *et al*., 2017). Activation of the proteases Caspase 1 (gene CASP1) (canonical) and Caspase 11 (gene CASP4) (non-canonical) represent the central outcomes of inflammasome activation, with VACV stimulation of the inflammasome well described (Amsler *et al*., 2013). ALVAC is also reported strongly to stimulate the inflammasome via AIM2 in both human and mouse cells (Liu *et al*., 2017). Viron assembly is arrested at the viroplasma stage in SCV-infected host cells due to the deletion of D13L (Eldi *et al*., 2017), which may limit inflammasome activation.

Poxviruses encode a number of proteins that seek to limit the activity of host immune responses (S1B Table, yellow highlighting), with some of these inhibiting the activities of cytoplasmic sensors (Fig. 2B, blue text). VACV’s decapping enzymes (D9 and D10, encoded by D9R and D10R) are expressed at the vaccination site (S1B Table). Both proteins inhibit dsRNA accumulation, and D10 is functional in mice (Liu *et al*., 2015); whether D9 is functional in mice is unknown (Fig. 2B, green cross). DHX9 is involved in both DNA and RNA sensing and is targeted by VACV’s E3 protein (encoded by E3L), with PKR inhibition the best defined activity of E3 (Brandt *et al*., 2001; Dempsey *et al*., 2018; Langland *et al*., 2006). VACV DNA is usually shielded from cytoplasmic sensors during replication in viral factories via wrapping in ER membranes; however, this wrapping is lost during virion assembly (Tolonen *et al*., 2001). The DNA sensor PRKDC/DNA-PK has a z-score of zero, perhaps due to the inhibitory activity of C4 (encoded by C4L) (Scutts *et al*., 2018), with C16L transcripts not detected by RNA-Seq in SCV12hQ (Fig. 2B, red cross; S1B Table). CrmA (from cowpox) and VACV’s homologue, B13R/SPI-2, inhibit caspase 1 (and other caspases), but are not functional in the Copenhagen strain of VACV (Smith *et al*., 2013) (Fig. 2B, black cross). NLRP1 was not identified by the IPA USR analysis, potentially due to the expression of F1L (S1B Table) (Gerlic *et al*., 2013).

### Dominant TLR-signaling associated signatures at 12 hours post vaccination

Following stimulation of TLR4, 9, 7 and 2 (but not TLR3) (Fig. 2A), a series of signaling events are initiated via the Myddosome, which contains MyD88/IRAK2/IRAK4 and signals to IRAK1 and TRAF6, with TRAF3 acting as a negative regulator (Kawasaki *et al*., 2014; O’Neill *et al*., 2010). TRAF5 (Buchta *et al*., 2014), TRAFD1/FLN29 (Sanada *et al*., 2008) and IRAK3 (aka IRAK-M) (Kobayashi *et al*., 2002) are also negative regulators of TLR signaling. TLR3 signaling also involves TRAF6 and TRAF3. All the aforementioned signaling molecules were identified by the IPA USR analysis, with negative regulators having negative z-scores and the rest positive z-scores (Fig. 3A; S2D Table, direct and indirect). These results are entirely consistent with the dominant TLR signatures illustrated in Fig. 2A. IL-1 receptor signaling (see below) also involves many of the same signaling molecules as TLR signaling (Rhyasen *et al*., 2015). The dominance of TBK1 may reflect its involvement in a series of signaling pathways; specifically, TLRs (including TLR3), STING and MDA5/MAVS (Zhao *et al*., 2019) that were illustrated in Fig. 2. C6 (encoded by C6L) inhibits TBK1 via binding to TBK1 adaptors (such as TANK) (Fig. 3A) thereby inhibiting activation of IRF3 and IRF7 (Smith *et al*., 2018; Unterholzner *et al*., 2011). K7 (encoded by K7R) binds DDX3 (Oda *et al*., 2009), an adaptor protein for the TBK1/IKKε complex that promotes IRF3 phosphorylation (García-Arriaza *et al*., 2013; Smith *et al*., 2018). TLR signaling is also inhibited by K7 and A52 (encoded by A52R), which bind to IRAK2 and TRAF6 (Carty *et al*., 2010; Smith *et al*., 2013; Yokota *et al*., 2010). N1 (encoded by N1L) binds TBK1 and inhibits NF-κB and IRF3 signaling pathways (Dai *et al*., 2014; DiPerna *et al*., 2004; Maluquer de Motes *et al*., 2011) (Fig. 3A).

**Figure 3.**
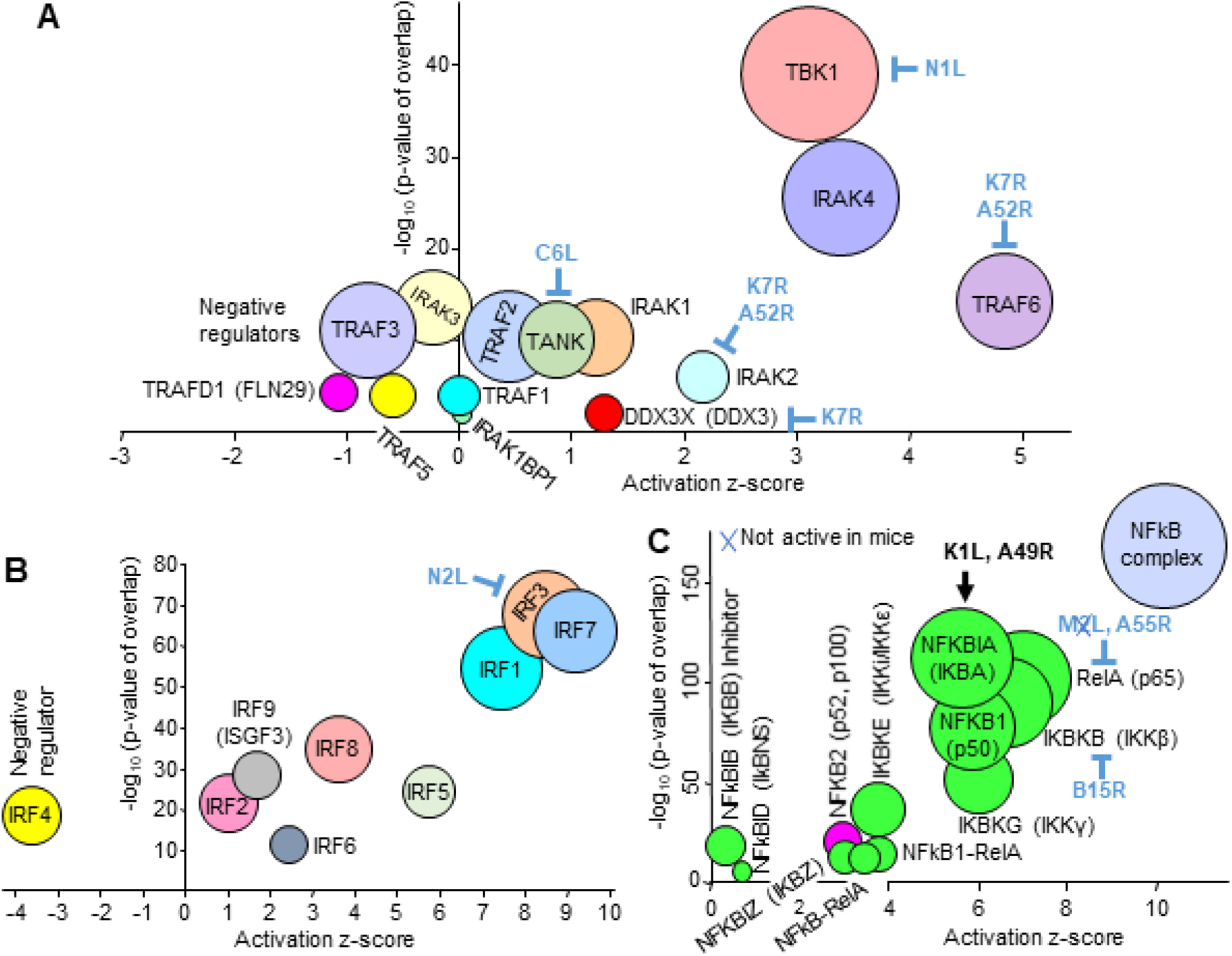
Secondary messenger signatures at 12 hours post vaccination. (A) Secondary messenger signatures. VACV encodes a series of cytoplasmic inhibitors, which are indicated in blue text. (B) Interferon response factors (IRFs). (C) NF-κB signatures. Green fill indicates canonical pathways, magenta fill non-canonical pathway, blue fill - not assigned to canonical or non-canonical. VACV-encoded cytoplasmic inhibitors are shown in blue text. K1L and A49R (black text) enhance the activity of NFKBIA, an inhibitor of the canonical pathway. Blue cross means the inhibitor is not active in mice.

### Interferon response factor signatures at 12 hours post vaccination

Interferon response factors (IRFs) are key transcription factors triggered by PAMPs, with IRF3, IRF7 and IRF1 dominating (Fig. 3B) in the IPA USR analysis (S2D Table, direct and indirect) of the DEGs from MQ vs SCV12hQ (S2B Table). These 3 IRFs are also in the top 5 USRs (sorted by p value) when the “direct” only option was used for the IPA USR analysis (S2D Table, direct only).

IRF3 and IRF7 are activated via multiple PAMP sensors described in Fig. 2 (Bakshi *et al*., 2017; Pham *et al*., 2016; Zhao *et al*., 2019), are intimately involved in driving antiviral responses (Jefferies, 2019; Rudd *et al*., 2012; Wilson *et al*., 2017), are also activated by MVA (Dai *et al*., 2014) and ALVAC (Harenberg *et al*., 2008), and have been shown to promote adaptive immune responses in a number of settings (Hatesuer *et al*., 2017; James *et al*., 2018; Suschak *et al*., 2016). SCV also encodes N2L, with N2 inhibiting IRF3 activation (Smith *et al*., 2013). IRF1 has a role in positive feedback maintenance of ISG expression (Michalska *et al*., 2018; Wilson *et al*., 2017) and has been shown to promote adaptive immunity in certain settings (Forero *et al*., 2019; Yang *et al*., 2018). IRF8 has a critical role in development and maturation of myeloid cells such as dendritic cells (Salem *et al*., 2020) and IRF5 is predominantly expressed by myeloid cells and regulates inflammatory responses, generally downstream of TLR-MyD88 pathways (Forbester *et al*., 2020). IRF4 is a negative regulator of TLR signaling (Negishi *et al*., 2005) (Fig. 3B).

### Conical NF-κB family signatures at 12 hours post vaccination

The NF-κB family of transcription factors play key roles in immunity, with the IPA USR analysis (S2D Table, direct and indirect) illustrating a dominant canonical NF-κB signature (Fig. 3C, green circles), consistent with the TLR signaling USRs described above. The dominance of the NFKBIA, but not another NF-κB inhibitor NFkBIB, likely reflects the activities of the K1L and A49R genes; both K1 and A49 proteins prevent degradation of NFKBIA (Smith *et al*., 2013), with A49 binding the ubiquitin ligase B-TrCP (Neidel *et al*., 2019). B14 (encoded by B15R in the Copenhagen strain) binds and inhibits IKBKB (Tang *et al*., 2018), and intracellular M2 (encoded by M2L) inhibits RelA (p65) nuclear translocation, but is not active in mice (Smith *et al*., 2013) (Fig. 3C, blue cross). A55 (encoded by A55R) dysregulates NF-κB signaling by disrupting p65-importin interaction, is active in mice (Pallett *et al*., 2019) and is expressed by SCV at the injection site (S1B Table).

### Th1 cytokine signatures at 12 hours post vaccination

The cytokine USR profile at 12 hours post vaccination is dominated by cytokine signatures generally associated with Th1 responses (Fig. 4A, red circles), in particular TNF, IL-1β and IFNγ, with *in vivo* induction of these cytokines by VACV suggested by previous studies (Carpenter *et al*., 1994; Tian *et al*., 2012; Tian *et al*., 2017). The Th1 dominance is consistent with studies on recombinant MVA vaccines (Bohnen *et al*., 2013; Tameris *et al*., 2014). TNF is required for optimal adaptive immune responses to VACV and other immunogens (So *et al*., 2019; Tian *et al*., 2017). IL-1 is important for host immune responses to VACV (Tian *et al*., 2012), with many vaccine adjuvants also inducing the release of IL-1 (Munoz-Wolf *et al*., 2018). Finally, IFNγ has anti-VACV activity (Kohonen-Corish *et al*., 1990) and has adjuvant properties in a range of settings (Nimal *et al*., 2005; Playfair *et al*., 1987; van Slooten *et al*., 2000). Although IL-27 was initially associated with Th1 responses, it is now recognized as a promoter of T regulatory cells (Yoshida *et al*., 2015) (Fig. 4A, IL27).

**Figure 4.**
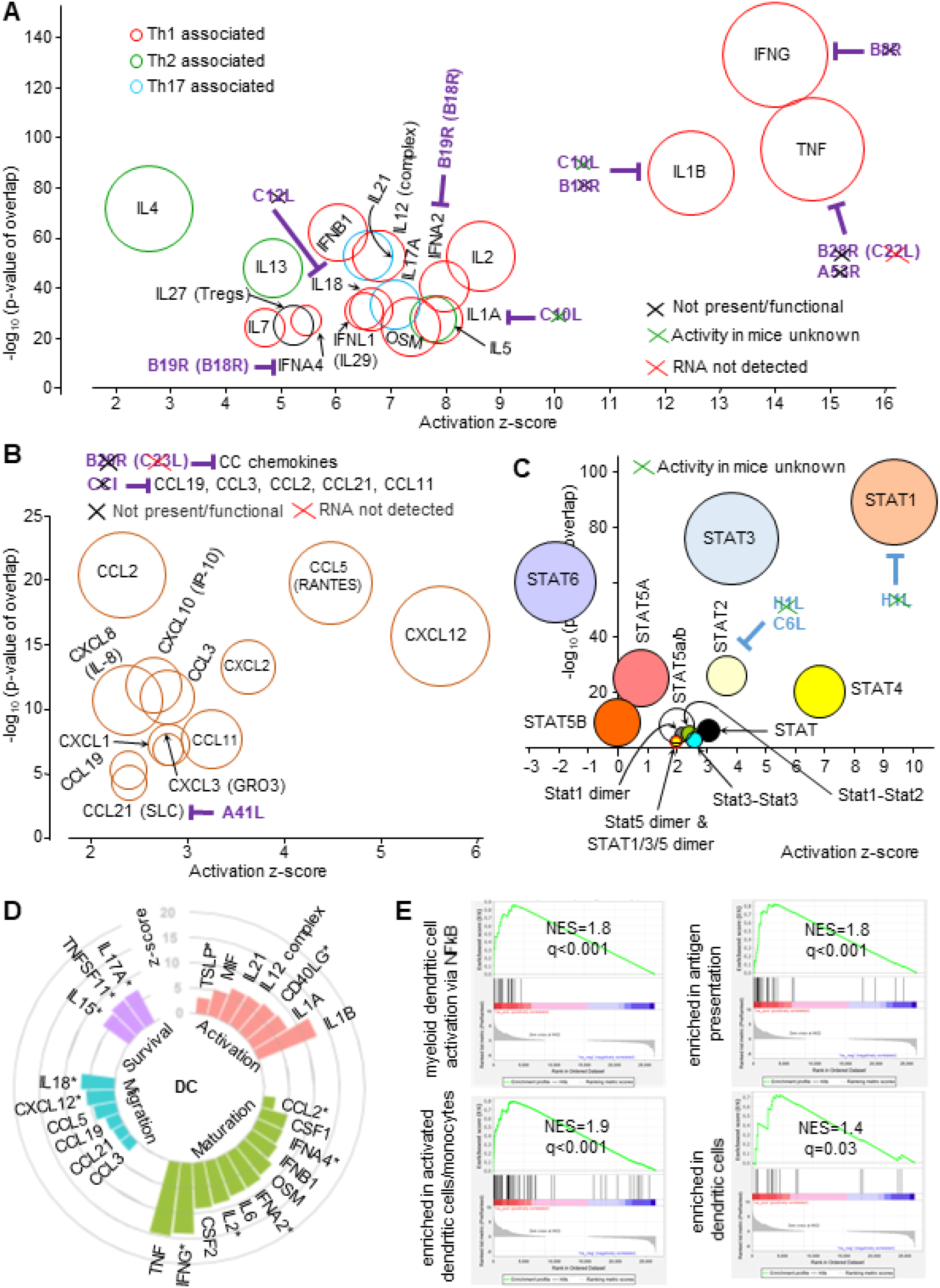
Cytokine, chemokine, dendritic cell and STAT signatures at 12 hours post vaccination. (A) Cytokine signatures. VACV genes encoding secreted inhibitors are shown in purple text. Black crosses indicate the inhibitors are not present or functional in SCV. Green crosses indicate that the activity in mice is unknown. Red crosses indicate the RNA was not detected in our RNA-Seq analysis. (B) Chemokine signatures. Purple text and crosses as in A. (C) STAT signatures. VACV encoded cytoplasmic inhibitors are shown in blue text. Crosses as in A. (D) IPA USRs associated with stimulation of dendritic cells. * indicates that the mediators have more than one of the four dendritic activities indicated. (The figure includes some USRs present in previous bubble graphs). For references see S2F Table. (E) GSEAs for the Blood Transcript Modules (right to left) M43.0 and M43.1 (gene sets combined, n=21); M95.0, M95.1, M71 and M200 (gene sets combined, n=49); M64, M67, M119 and M165) (gene sets combined, n=71); M168 (n=19). For gene set details see S2F Table.

VACV encodes a number of soluble inhibitors of several cytokines (Fig. 4A, purple text). C10L encodes C10, which blocks interaction of IL-1 with its receptor (Kluczyk *et al*., 2002), but its activity in mice is unknown (Fig. 4A, green cross). B16 (encoded by B16R) (B15R in Western Reserve) is a secreted IL-1β decoy receptor (Alcami *et al*., 1992; Perdiguero *et al*., 2009), but appears to be truncated in the Copenhagen strain of VACV (Uniprot; GCA 006458465.1). B8 (encoded by B8R) is a secreted IFNγ receptor homologue, and ZIKA prME was inserted into the B7R-B8R locus in SCV-ZIKA/CHIK thereby inactivating these genes (Prow *et al*., 2018b). B28R (C22L) and A53R encode TNF receptor homologues that are not active in the Copenhagen strain of VACV (Alcam *et al*., 1999). C12 (encoded by C12L) inhibits IL-18 (Symons *et al*., 2002). B19 (encoded by B19R) (also known as B18R in other VACV strains) is a decoy receptor for soluble IFNαs (Colamonici *et al*., 1995; Waibler *et al*., 2009). B19 mRNA is well expressed at the injection site (S1B Table) and is potentially responsible for the relatively low z-scores of the IFNα USRs (Fig. 4A). A35 (encoded by A35R) is an intracellular VACV protein that also inhibits the synthesis of a number of chemokines and cytokines (including IFNα, MIP1α, IL-1β, IL-1α, GM-CSF, IL-2, IL-17, GRO1/KC/CXCL1, RANTES, TNFα) by VACV-infected cells (Rehm *et al*., 2010) (not shown).

### Chemokine signatures at 12 hours post vaccination

The chemokine signatures (Fig. 4B) are dominated by (i) CXCL12, which is made by many cell types and is strongly chemotactic for lymphocytes, (ii) CCL5 (RANTES), which is *inter alia* chemotactic for T cells and (iii) CXCL2, a neutrophil chemoattractant (consistent with Fig. 1E). CXCL12 and CCL5 are also involved in dendritic cell (DC) recruitment (Chabot *et al*., 2006; Lopez *et al*., 2018). CCL2 is also induced by MVA (Lehmann *et al*., 2009; Lehmann *et al*., 2016) and is involved in DC maturation and induction of T cell immunity (Gomes *et al*., 2019).

VACV infected cells secrete a number of proteins that bind and inhibit certain chemokines, although only A41 (encoded A41L) is active in the Copenhagen strain of vaccinia (Fig. 4B, purple text). A41L inhibits CCL21 (Smith *et al*., 2013), perhaps consistent with its low z-score (Fig. 4B). B29, encoded by B29R (C23L) inhibits multiple CC chemokines, but is inactive in the Copenhagen strain (Alcamí *et al*., 1998). CCI inhibits a series of chemokines (Fig. 4B), but is not expressed on the Copenhagen strain of VACV (Burns *et al*., 2002; Reading *et al*., 2003).

### STAT signatures at 12 hours post vaccination

Cytokines and chemokines bind to their receptors and activate transcription via STATs. The dominant STAT signatures (from S2D Table, IPA direct and indirect) were STAT1, STAT4 and STAT3 (Fig. 4C), with STAT1 and STAT3 representing the top USRs by p value and STAT1 also the top USR by z-score when analyzed by IPA using direct only interaction (S2D Table, direct only). STAT1 forms complexes primarily STAT1-STAT1 homodimers (stimulated by IFNγ signaling) and STAT1-STAT2-IRF9 (ISGF3) (stimulated by type I IFN signaling). Using Interferome to interrogate the “Target molecules in dataset” listed for the STAT1 signature (S2D Table, direct and indirect), nearly all the target molecules were deemed IFNγ inducible (with most also inducible by type I IFNs). The relatively low p values and z-scores for STAT1 dimers would thus appear to be an under-annotation within IPA. The dominant STAT1 signature (Fig. 4C) is thus consistent with the dominant IFNγ signature in Fig. 4A. The cytoplasmic VACV-expressed H1 (encoded by H1L) inhibits STAT1 and STAT2 (Mann *et al*., 2008), but clearly only in cells infected with VACV. C6 (encoded by C6L), as well as the aforementioned binding of TBK1 adaptors, also binds the TAD domain of STAT2 (Stuart *et al*., 2016).

STAT4 signaling is induced by a number of cytokines including IL-12 and IL-2 (Yang *et al*., 2020) and is critical for IFNγ production during generation of Th1 responses (Varikuti *et al*., 2016). STAT3 signaling is induced by a number of cytokines including IL-6 and OSM (and growth factors such as GM-CSF), with BCG vaccination recently shown to cause STAT3 phosphorylation in antigen presenting cells (Copland *et al*., 2019). STAT6 is involved in driving Th2 responses (Gaylo-Moynihan *et al*., 2019) and has a negative z-score (Fig. 4C), consistent with the Th1 dominance illustrated in Fig. 4A.

### Dendritic cell associated signatures

A range of mediators affect dendritic cell activities, with many of these identified as USRs by IPA analysis of DEGs for MQ vs SCV12hQ (Fig. 4D; S2D Table; for references see S2F Table). The VACV protein A35 inhibits a number of these mediators (see above), as well as inhibiting class II antigen presentation (Rehm *et al*., 2010). Multiple key mediators needed for induction of adaptive immune responses by dendritic cells would thus appear to be active at the injection site 12 hours post vaccination.

Extensive bioinformatics treatments of >30,000 peripheral blood transcriptomes from >500 human studies of 5 vaccines provided 334 publically available gene sets in the form of Blood Transcription Modules (BTMs). BTM gene sets are associated with specific subsets of cells and/or their activities (Li *et al*., 2014). BTM gene sets associated with dendritic cells and dendritic cell activities (S2F Table) and Gene Set Enrichment Analyses (GSEAs) were used to determine whether genes from dendritic cell BTMs were significantly represented in the MQ vs SCV12hQ gene list (S2A Table). The GSEAs provided highly significant results (Fig. 4E), illustrating that signatures associated with dendritic cells and their activities can be readily identified at the injection site 12 hours post vaccination. Such signatures likely underpin the immunogenicity of the vector system.

### Injection site signatures at day 7 post vaccination

The most common side effects reported for MVA (licensed as a small pox vaccine in Europe as IMVANEX) were at the site of subcutaneous injection; most of them were mild to moderate in nature and resolved without any treatment within seven days. To gain insights into the injection site responses after SCV vaccination, RNA-Seq of muscles on day 7 post-vaccination was undertaken to provide a gene list (MQ vs SCVd7Q, S2G Table), from which a DEG list (n=1413 genes) was generated (S2H Table) by applying the same filters as above (q ≤0.01, FC ≥2 and sum of all counts across the six samples >6). Of the 1413 DEGs, 1337 were up-regulated, with 633 (47%) of these also up-regulated DEGs for MQ vs SCV12hQ.

Cytoscape analyses of up-regulated DEGs from MQ vs SCV12hQ (S2C Table) were compared with MQ vs SCVd7Q (S2I Table). Multiple top signatures (by FDR) associated with T cells and B cells were substantially more significant on day 7 than at 12 hours (Fig. 5A; Table S2J). For instance, FDR values associated with the GO Process terms “positive regulation of T cell activation” and “T cell differentiation” were ≈9 logs more significant by day 7, when compared with 12 hours post vaccination (Fig. 5A; S2J Table). T cell receptor associated KEGG Pathways and GO Component terms were also more significant on day 7 (Fig. 5A; S2J Table). “T cell receptor complex” was also the top “GO Cellular Component” term by p value for day 7 up-regulated DEGs (S2J Table, Enrichr). A similar pattern emerged for B cell terms (Fig. 5A; S2J Table).

**Figure 5.**
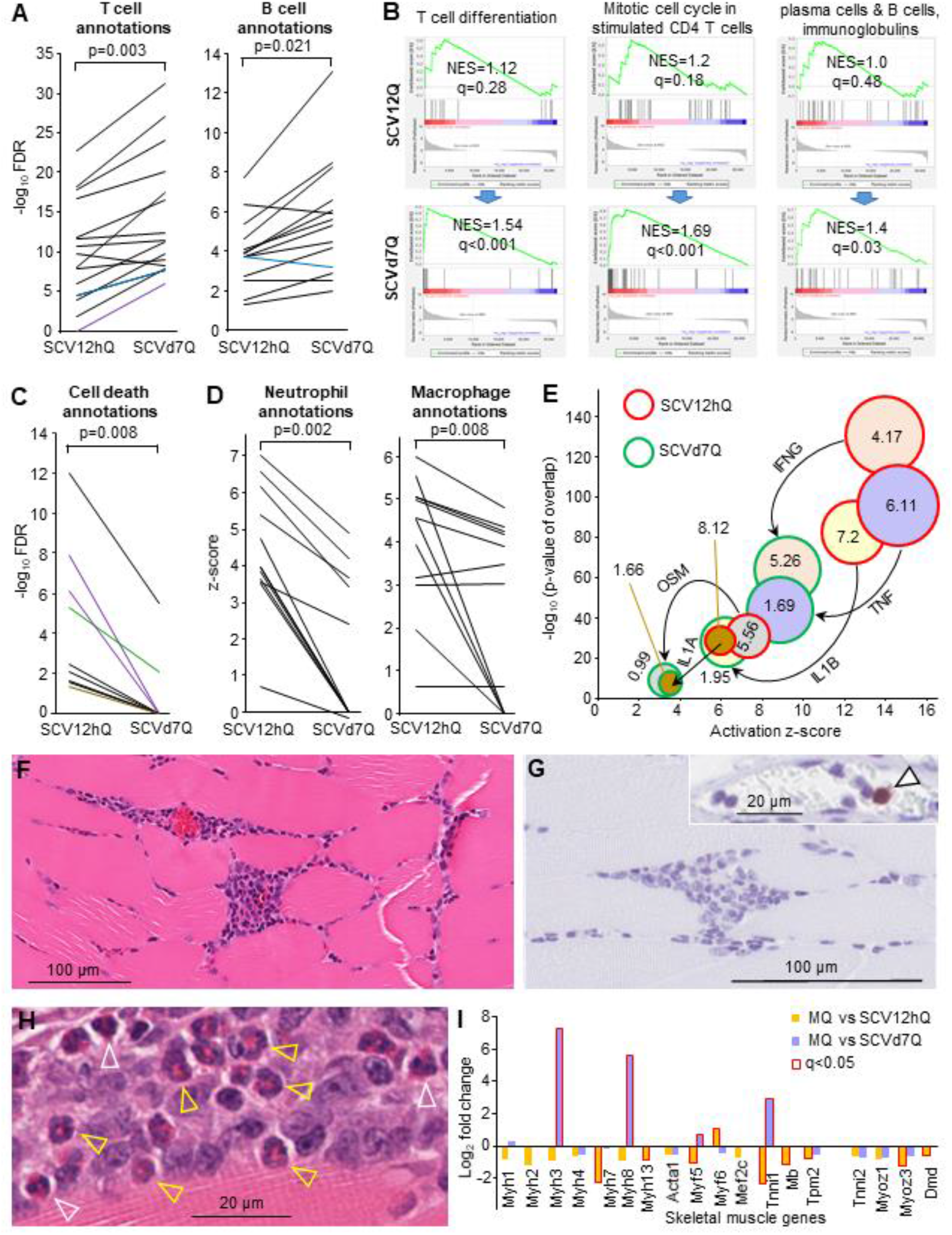
The injection site day 7 post vaccination. (A) Cytoscape analysis of up-regulated DEGs from MQ vs SCV12hQ and MQ vs SCVd7Q, illustrating the upward trend in significance of top B and T cell associated annotations. Black line – GO Process, Blue line GO - Component, Purple line - KEGG Pathways. For full lists and descriptions of annotations see S2J Table; statistics by paired t test for the full lists (parametric data distribution). (B) GSEAs use to interrogate MQ vs SCV12hQ and MQ vs SCVd7Q gene lists using T cell and B cell BTMs (left to right) M14 (n=12); M4.5 (n=35); M156.0 and M156.1 (gene sets combined, n=56) (for details of BTM gene sets see S2J Table). (C) As for A illustrating the downward trend of cell death annotations. Anotations not identified by the IPA analysis were nominally given a −log_10_ FDR value of zero (y axis). Color coding as for A, but also Green line – UniProt Keywords, Brown line – Reactome Pathways. For descriptions of annotations see S2K Table. (Statistics by Wilcoxon Signed Rank tests; non-parametric data distribution). (D) IPA *Diseases and Functions* analysis of DEGs (up and down-regulated) from MQ vs SCV12hQ and MQ vs SCVd7Q, illustrating the downward trend in z-scores for macrophage and neutrophil annotations (for description of annotations see Table S2L and S2M). (Statistics by Wilcoxon Signed Rank tests). (E) Major IPA USR pro-inflammatory cytokine annotations identified at 12 hours (Fig. 4A; S2D Table) had much lower z-scores and p values on day 7 post vaccination (S2N Table). Numbers in the circles represent the log_2_ fold change for that cytokine relative to MQ. (F) H&E staining of intramuscular injection site lesions on day 7 post infection. (G) Neutrophil Ly6G staining of lesions from day 7 post infection. Arrow in insert shows positive staining of a neutrophil in a blood vessel capillary. (H) Eosinophils in the intramuscular injection site lesions on day 7 post vaccination. White arrow heads - mature segmented eosinophils. Yellow arrowheads – immature band eosinophils. (I) Expression of skeletal muscle genes from MQ vs SCV12hQ (S2a Table) and MQ vs SCVd7Q (S2G Table); bars with red outline indicate significant fold change (q<0.05).

GSEAs (as in Fig. 4E) using genes from BTMs (Li *et al*., 2014) associated with T cell differentiation and division, and B cell differentiation into plasma cells, showed significance for SCVd7Q, but not SCV12hQ (Fig. 5B). Thus remarkably, these BTMs were able to identify signatures at the injection site on day 7 that were associated with the development of adaptive immune responses. (IgG responses are known to be induced after SCV-ZIKA/CHIK vaccination (Prow *et al*., 2018b)).

Vaccination site lesions are well described for VACV vaccination (Frey *et al*., 2002), with skin lesions reported days 6-11 after vaccination with the Lister strain (Talbot *et al*., 2006) and days 3-19 after Dryvax vaccination (Parrino *et al*., 2007). Such lesions are associated with cell death (He *et al*., 2014), tissue damage (Fulginiti *et al*., 2003) and recruitment of neutrophils, with neutrophil recruitment also a feature of *eczema vaccinatum*, a complication of smallpox vaccination (Darling *et al*., 2014; Frey *et al*., 2002; Tian *et al*., 2017). Cytoscape analyses (S2C and S2I Table) illustrated that the cell death pathway annotations identified at 12 hours (Fig. 1G) were considerably less significant or absent for day 7 (Fig. 5C, S2K Table). Analysis of the 1413 DEGs from MQ vs SCVd7Q (S2H Table) with IPA *Diseases and Functions* feature (S2L Table), showed a significant reduction in the z-scores of neutrophil-associated annotations on day 7 when compared to 12 hours (Fig. 5D; S2M Table). (The Cytoscape analysis also showed a highly significant reduction in FDR values for neutrophil terms, S2I Table, graph on right). These analyses indicate that progression of cell death and neutrophil infiltration is not a feature of SCV vaccination, likely consistent with the inability of SCV to produce viral progeny (Eldi *et al*., 2017). SCV does not cause a spreading infection, with vaccine-derived mRNA lost by day 7 (Fig. 1A). A similar IPA *Diseases and Functions* analysis of macrophage-associated annotations also illustrated a significant reduction by z-scores (Fig. 5D, S2M Table), further indicating that injection site inflammatory responses were abating by day 7 (Soehnlein *et al*., 2010).

The dominant pro-inflammatory cytokine USRs identified at 12 hours post vaccination (Fig. 4A) were substantially lower by day 7 post vaccination with respect to both −log_10_ p values and z-scores (Fig. 5E; S2D vs S2N Table). Fold changes in cytokine mRNA expression levels relative to MQ were also substantially lower on day 7 (Fig. 5E), with the exception of IFNγ, which had a fold change relative to MQ of 4.17 at 12 h and a fold change of 5.26 relative to MQ on day 7, perhaps due to the emerging Th1 T cell responses (see above). IPA *Diseases and Functions* also showed reduced significance and z scores on day 7 for *Inflammatory response* (−log_10_ p value 110.7 to 59.8, z-score 7.7 to 5.5) and *Chronic inflammatory disorder* (−log_10_ p value 70.1 to 38.1, z-score −0.23 to −2.2) (S2L Table). These analyses again argue that inflammation at the injection site is abating on day 7, with persistent inflammation at the injection site generally deemed undesirable in most vaccination settings (Clarke *et al*., 2020; Lee *et al*., 2008; Wang *et al*., 2014).

### Loss of neutrophils and presence of eosinophils on day 7 post vaccination

H&E staining of the injection sites day 7 post vaccination supports the bioinformatics results described in the previous section. When compared with 12 hours (Fig. 1D), necrotic muscle lesions were largely absent, with the cellular infiltrates less disseminated and more focal (Fig. 5F). In addition, in contrast to 12 hours (Fig. 1E), neutrophils (stained with anti-Ly6G) were not observed in the day 7 cellular infiltrates (Fig. 5G), although the occasional neutrophil could be seen in blood vessels, illustrating that the staining had worked (Fig. 5G, insert, arrowhead). In contrast to Fig. 1E, Apoptag staining was also largely negative on day 7 (not shown). Loss of neutrophils is consistent with inflammation resolution (Soehnlein *et al*., 2010).

Another feature of the resolving infiltrates on day 7 post vaccination (clearly evident from H&E staining) was the presence of eosinophils (Fig. 5H), despite the retention of a dominant Th1 signature (Fig. 5E). Many of these cells showed the morphological features of immature band eosinophils, as distinct from segmented mature eosinophils (Fig. 5H).

At 12 hours post vaccination, genes specific to skeletal muscle were generally slightly down-regulated (Fig. 5I; S2A Table), consistent with the SCV infection-associated necrosis or pyroptosis (Fig. 1D). On day 7 post vaccination, Myh3 and Myh8 were significantly up-regulated (Fig. 5I; S2G Table), with these genes transiently up-regulated after muscle injury (Yoshimoto *et al*., 2020). Tnni1, a skeletal myogenesis marker (Park *et al*., 2016), was also up-regulated (Fig. 5I). However, stable expression of Tnni2, Myoz1, Myoz3 and Dmd by day 7 (Fig. 5I), argues that muscle regeneration had been largely completed at this time (Yoshimoto *et al*., 2020); consistent with the H&E staining (Fig. 5F).

### No compelling arthritic signature in feet on day 7 post vaccination

An early attenuated CHIKV vaccine tested in a phase II human clinical trial caused a transient arthralgia in 5 out of 58 vaccine recipients (Edelman *et al*., 2000). Using RNA-Seq we have previously characterized the molecular signatures associated with hind foot arthritis 7 days after infection of mice with CHIKV (Prow *et al*., 2019; Wilson *et al*., 2017). Many features identified in mice recapitulated those seen in human patients (Michlmayr *et al*., 2018; Suhrbier, 2019; Wilson *et al*., 2017). We re-analyzed the FASTQ files (Wilson *et al*., 2017) (deposited in NCBI BioProject PRJNA431476) using STAR aligner and the more recent mouse genome build (GRCm38 Gencode vM23). The complete gene list for day 7 feet (peak CHIKV arthritis) is provided in S2O Table.

To determine whether SCV-ZIKA/CHIK vaccination is associated with the induction of an arthritic signature on day 7 post vaccination, feet were collected (by severing at the bottom of the tibias after euthanasia) and analyzed by RNA-Seq (MF vs SCVd7F) (S2P Table). A DEG list was generated after application of two filters q<0.05 and the sum of counts across all 6 samples >6, resulting in only 22 genes, of which 8 were up-regulated (S2Q Table). Three of these genes (Daglb, Tgtp2 and Gbp3) were also present in the up-regulated DEGs for day 7 feet of CHIKV infected mice (S2O Table), although the fold change of the latter two were substantially higher after CHIKV infection than after SCV vaccination (Fig. 6A). Of the 8 up-regulated DEGs, Tnfaip6 and Crispld2 have anti-inflammatory activities (Day *et al*., 2019; Zhang *et al*., 2016) and Thbs4 and Daglb have pro-inflammatory activities (Hsu *et al*., 2012; Rahman *et al*., 2020), with Gbp3 and Spon2 associated with antiviral responses (Li *et al*., 2009b; Nair *et al*., 2017) and Mrgprf associated with the itch response (Meixiong *et al*., 2017).

**Figure 6.**
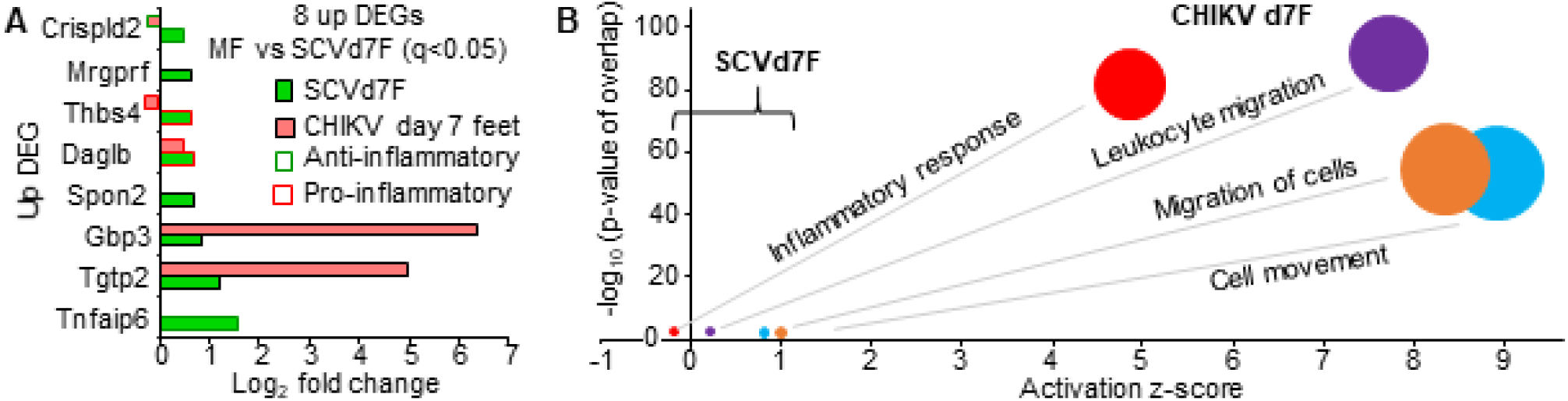
No compelling arthritic signature after SCV vaccination. (A) MF vs SCVd7F provided 22 DEGs (with 2 filters applied q<0.05 and count sum >6) of which 8 were up-regulated (Table S2Q). Three of these were also up-regulated DEGs for CHIKV arthritis day 7 post infection (Table S2O; q<0.05). (B) The 22 DEGs analyzed by IPA *Diseases and Functions* feature (direct and indirect) (S2Q Table) and compared with the same annotations identified by IPA analysis of DEGs for CHIKV arthritis (S2O Table).

IPA *Diseases and Functions* analysis of the 22 DEGs returned a significant “Inflammatory response” with a negative z-score, and cellular infiltrate terms with low z-scores (Table S2Q). Importantly, the z-scores and p-values for these annotations were very much lower for SCVd7F than they were for day 7 feet from CHIKV infected mice (Fig. 6B). Comparison of the 22 DEGs with those reported for collagen induced arthritis (CIA) (GSE13071) (Geurts *et al*., 2009) also identified no obvious concordance (S2Q Table), although with only 22 DEGs such a comparison is somewhat underpowered.

Overall these results suggest that SCV vaccination was not associated with a compelling arthritic signature, even though the injection sites (quadriceps muscles) were in the same legs as the feet that were used to generate the MF vs SCVd7F gene set.

## Discussion

We provide herein a detailed systems vaccinology analysis of a recombinant SCV vaccine in mouse muscle, to provide insights into vaccine gene expression, and host adjuvant signatures and immune responses. Of all the reads mapping to the SCV-ZIKA/CHIK vaccine, ≈20% mapped to the recombinant immunogens. IHC illustrated immunogen protein expression in skeletal muscle cells, with these cells showing histological signs of necrosis (Lentscher *et al*., 2020; Szugye, 2020) and bioinformatics indicating the presence of necrosis and necroptosis (Morgan *et al*., 2018). Adjuvant signatures were driven by TLRs, cytoplasmic RNA and DNA sensors and the inflammasome, with neutrophils potentially also contributing (Stephen *et al*., 2017). By day 7 vaccine transcripts and neutrophils were largely absent, and inflammation was abating, with the presence of what appeared to be tissue repair-associated eosinophils (Coden *et al*., 2020; Heredia *et al*., 2013; Lloyd *et al*., 2018; Toor, 2019; Weller *et al*., 2017). Although a previous live-attenuated CHIKV vaccine was associated with some arthropathy (Edelman *et al*., 2000), no compelling arthritic signature was evident after SCV vaccination.

The immunogenicity of SCV, and likely poxvirus vector systems generally, would appear to be underpinned by the ability to stimulate a broad range of adjuvant pathways. Multiple TLR signatures were identified at 12 hours post vaccination (Fig. 2A), with the TLR4 agonist monophosphoryl lipid A a component of Fendrix (hepatitis B vaccine) and Cervarix (human papilloma virus vaccine) (which are formulated in ASO4 adjuvant), as well as Shingrix (herpes zoster vaccine) (formulated in AS01 adjuvant). The dsRNA TLR3 agonists, Ampligen (Rintatolimod) and Hiltonol, also have well described adjuvant properties (Martins *et al*., 2015), with Hiltonol being tested in therapeutic cancer vaccine trials (ClinicalTrials.gov Identifier: NCT04345705 and NCT02423863). The TLR9 agonist, CpG oligonucleotide (CpG 1018), was recently approved as an adjuvant in Heplisav-B (hepatitis B vaccine). A range of cytoplasmic sensor signatures with known adjuvant activity were also identified (Fig. 2B), including multiple inflammasome signatures; the best known human adjuvant, alum, is believed to mediate its activity via activation of the inflammasome (Kelley *et al*., 2019). Stimulation of cytoplasmic dsRNA sensors represents a key adjuvant activity for replicon-based RNA vaccines (Fuller *et al*., 2020; Pijlman *et al*., 2006) and the utility of STING-activating adjuvants is being actively explored (Jiang *et al*., 2020; Luo *et al*., 2019). Although SCV does not generate viral progeny, it does replicate its DNA (Eldi *et al*., 2017), perhaps explaining the dominant STING signature (Fig. 2B). Virulent poxviruses inhibit STING activation via unknown factors, an inhibitory activity not found for MVA (Georgana *et al*., 2018). This activity is perhaps similarly absent for SCV or is inactive in mice.

A desirable feature for any vaccine is the avoidance of reactogenicity, a term describing a series of post-vaccination adverse events often associated with excessive injection site inflammation and systemic reactions such as fever (Herve *et al*., 2019). A potential goal of systems vaccinology is the identification of reactogenic signatures; however, consensus regarding the composition of such signatures, and/or when and where best to sample to obtain such signatures, has yet to be established (Gonzalez-Dias *et al*., 2020). CCL2 and CXCL10 up-regulation in peripheral blood was identified as potential biomarkers of vaccine-elicited adverse inflammation in mice after a number of different vaccines given i.m. (McKay *et al*., 2019). These chemokines featured prominently at the injection site 12 hours after SCV vaccination (Fig. 4B), although fold-change had reduced substantially by day 7 (log_2_ 4.12 to −0.73, and 8.35 to 3.17, respectively) (S2A and S2G Table). CCL2 is also induced by MVA (Lehmann *et al*., 2016), can be important for avoiding immunopathology (Poo *et al*., 2014), and is induced by the licensed adjuvants, Alum and MF59 (Seubert *et al*., 2008). CXCL10 is also induced by MVA (Flechsig *et al*., 2011) and by the licensed adjuvant, MF59 (Schifanella *et al*., 2019). Given the extensive clinical safety record of MVA vaccination (Overton *et al*., 2018) and the lack of overt injection site reactogenicity or fever observed in NHPs after SCV vaccination (Prow *et al*., 2020), CCL2 and CXCL10 up-regulation at the injection site would thus not appear to be compelling biomarkers for adverse events after MVA or SCV vaccination. Similarly, a whole-blood systemic adverse event signature for yellow fever 17D vaccination has been reported, with 32 up-regulated genes on day 1 (but not day 3) associated with a range of systemic adverse events (either within 24 hours or a median time post vaccination of 6 days) (Chan *et al*., 2017). GSEAs illustrate that this signature is highly significantly present in the SCV12hQ gene list, and is also significantly present in the SCVd7Q gene list (S2R Table). All the core enriched genes (S2R Table) were type I IFN stimulated genes (by Interferome), with type I IFN stimulated gene induction clearly present at the SCV injection site (Fig. 4A). However, MVA vaccination is also associated with short term (i) increases in local type I IFN responses (Dai *et al*., 2014; Wolferstatter *et al*., 2014) and (ii) elevated serum IFNα levels (Lopez-Gil *et al*., 2013; Waibler *et al*., 2007). Of note, SCV encodes B19R/B18R, an inhibitor of type I IFN responses (Waibler *et al*., 2009). Unlike Yellow fever 17D, MVA and SCV vaccinations are not associated with significant viremias or viral dissemination, reducing the probability of excessive serum type I IFN responses and systemic adverse events; although pyrexia, headache, myalgia, nausea, fatigue and/or chills are seen in a small percentage of MVA vaccine recipients (Overton *et al*., 2018). Clearly, adverse event signatures identified in peripheral blood, may not to be overly informative for understanding adverse events at the injection site. In addition, not only the presence of specific gene transcripts but also the magnitude of gene induction are likely to be important, with the latter not fully taken into account by GSEAs. In humans, sampling injection sites is difficult, although emerging micro-sampling techniques may provide new avenues (Lei *et al*., 2019). Ultimately RNA-Seq of injection sites in animal models should be able to provide early warnings in the vaccine development process of potential reactogenicity issues. Herein we show that, although SCV (unlike MVA) retains the ability to replicate its DNA ((Eldi *et al*., 2017), the injection site reactogenicity (like IMVANEX) has largely resolved by day 7 post vaccination.

How might the information provided herein find utility for poxvirus vaccine design? The ZIKA and CHIK immunogens are inserted into B7R/B8R and A39R, respectively (Prow *et al*., 2018b). The 12 vector genes that were not expressed *in vivo* post-vaccination (S1B Table) offer other potential insertion sites for recombinant immunogens that would ostensibly have minimal impact on vaccine behavior. However, expression of these genes in human muscle might be checked, perhaps via use of human skeletal muscle organoids (Gholobova *et al*., 2019). The multiple adjuvant pathway stimulated by SCV (Fig. 2) might argue for a certain level of redundancy (Waibler *et al*., 2007), which might allow certain inhibitors to be reintroduced with the aim of reducing reactogenicity, without compromising immunogenicity. For instance, B13R (also known as SPI-2) is absent in the Copenhagen strain of VACV and in SCV, and inhibits caspase I, a key protease for generation of bioactive IL1-β (Martin-Sanchez *et al*., 2016). Reintroducing SPI-2, or B16R that encodes an IL-1 receptor (Jackson *et al*., 2005), into the vector might have minimal effects on immunogenicity (Legrand *et al*., 2004), whilst reducing generation of this potent pyrogen and thereby potentially the risk of adverse events such as fever (Alcami *et al*., 1992). Recent sequencing of ancient Variola viruses from Viking corpses perhaps supports such strategies, as active expression of immune modulating genes may be associated with reduced pathogenicity (Alcami, 2020; Muhlemann *et al*., 2020). Co-formulation of SCV with adjuvants (Barrera *et al*., 2018; Magnusson *et al*., 2018) or encoding genetic adjuvants within SCV (Matchett *et al*., 2020) might appear superfluous, given the large number of adjuvant pathways already being activated. Introducing apoptosis inhibitors to improve immunogenicity (Chea *et al*., 2019) may have minimal impact for i.m. injections of SCV (or other pox vectors) as skeletal muscle does not appear readily to undergo apoptosis (Schwartz, 2008, 2018), with B13R also well expressed at the injection site (S1B Table). The absence of the chemokine inhibitors B29R (C23L) (not expressed) and CCI (not present/functional) may contribute *inter alia* to effective recruitment of dendritic cells. Deletion of A41L might be tested to determine whether this would increase immunogenicity, given CCL21 recruits T cells and enhances T-cell responses (Gray *et al*., 2018). Complement control proteins VCP (encoded by B27R) and C3 (encoded by C3L) were expressed at the mRNA level, with VCP deletion from VACV increasing anti-VACV antibody responses (Albarnaz *et al*., 2018), suggesting deletion of these genes might enhance immunogenicity. One might consider deletion of N2L (an inhibitor of IRF3) as this was shown to improve the immunogenicity of a recombinant MVA vaccine (Garcia-Arriaza *et al*., 2014) and also reduced the virulence of VACV (Ferguson *et al*., 2013). However, the IRF3 signature is already very dominant (Fig. 3B), so additional IRF3 activation (if possible) may not translate to significant improvements in immunogenicity. Deletion of C6L increased the immunogenicity of a recombinant MVA vaccine (Marin *et al*., 2018), presumably by relieving IRF3, IRF7 (Smith *et al*., 2018; Unterholzner *et al*., 2011) and/or STAT2 inhibition (Stuart *et al*., 2016). However, C6L deletion could risk excessive type I IFN responses and increased reactogenicity (Chan *et al*., 2017).

The presence of eosinophils in the resolving lesion on day 7 was unexpected. Eosinophils have been reported in pruritic papulovesicular eruptions in a case of generalized vaccinia after smallpox vaccination (Beachkofsky *et al*., 2010) and are well described as drivers of allergic diseases such as eosinophilic asthma (Bhalla *et al*., 2020; Côté *et al*., 2020). However, recently a role for eosinophils in tissue repair and wound healing has emerged (Coden *et al*., 2020; Lloyd *et al*., 2018), particularly for muscle tissues (Heredia *et al*., 2013; Toor, 2019; Weller *et al*., 2017). That the eosinophils in these resolving post-vaccination lesions are distinct from inflammatory eosinophils is supported by the absence of IL-5 mRNA expression (S2G Table), with anti-IL-5 therapy used for eosinophilic asthma (Walsh, 2020). Transcripts for eosinophil cationic protein (Ear1) and eosinophil peroxidase (Epx) (granule components of inflammatory eosinophils) were also not detected. Eotaxins (CCL11, CCL24 and CCL26) were not up-regulated, with IL4, IL13, IL3 and GM-CSF (CSF2) transcripts absent (S2G Table). We were unable to find any eosinophil gene signatures that provided a significant result after GSEAs of the MQ vs SCVd7Q gene set, suggesting that signatures for inflammatory eosinophils are distinct from tissue repair-associated eosinophils, with the signatures for the latter yet to be defined.

A limitation of using mice to analyze VACV-based vaccines is that several VACV-encoded inhibitors are not active in mice (encoded by M2L, A38L) and the activity of others in mice is not known (D9R, H1L, C10L). The activity in mice of B28R and B29R is also not known, but these inhibitors are not active in the Copenhagen strain of VACV. Others were found not to be expressed in mice (C16L, C23L, C22L), although it is unclear whether they are poorly expressed generally or poorly expressed in muscle or poorly expressed in mice. How critical these genes are to the overall interpretations presented herein is difficult to assess, given the presence of multiple overlapping and potentially cross-compensating pathways.

An extensive history of poxvirus vector development has yet to culminate in the licensing of a recombinant poxvirus vector for human use, although several are in late stage clinical trials. Systems vaccinology approaches such as those described herein may facilitate rationale refinement of pox vector design and contribute to progressing such systems towards registration and licensure.

## Methods

### Ethics statement

All mouse work was conducted in accordance with the “Australian code for the care and use of animals for scientific purposes” as defined by the National Health and Medical Research Council of Australia. Mouse work was approved by the QIMR Berghofer Medical Research Institute animal ethics committee (P2235 A1606-618M).

### Mice and vaccination

Female C57BL/6J mice (6-8 weeks) were purchased from Animal Resources Center (Canning Vale, WA, Australia). Mice were vaccinated once with 50 μl of 0.5 x 10^6^ pfu SCV-ZIKA/CHIK i.m. or Mock vaccinated (with PBS) into both quadriceps muscles as described (Prow *et al*., 2018b).

### RNA-Seq

At the indicated times post vaccination, mice were euthanized using CO_2_ and quadriceps muscles or feet placed individually into RNAlater (Life Technologies) overnight at 4°C and then homogenized in TRIzol (Invitrogen) using 4 x 2.8 mm ceramic beads (MO BIO Inc., Carlsbad, USA) and a Precellys24 Tissue Homogeniser (Bertin Technologies, Montigny-le-Bretonneux, France) (6000 rpm on ice, 3 times 12 sec for feet and 2 times for 10 seconds for muscle). Homogenates were centrifuged (14,000 g x 15 min) and RNA extracted from the supernatants as per manufacturer’s instructions. Following DNase treatment (RNAseq-Free DNAse Set (Qiagen)) and RNA purification (RNeasy MinElute Kit), RNA concentration and purity was determined by Nanodrop ND 1000 (NanoDrop Technologies Inc.). RNA samples were pooled so that for each group of 6 mice, 12 sets of quadriceps muscles or feet were used to create 3 biological replicates which contained equal amounts of RNA from 4 different mice. All replicates were then sent to the Australian Genome Research Facility (Melbourne, Australia) for library preparation and sequencing.RNA integrity was assessed using the Bioanalyzer RNA 6000 Nano assay (Agilent) and libraries were prepared from 200 ng of total RNA using TruSeq Stranded mRNA library preparation kit (Illumina). The resulting libraries were assessed by TapeStation D1K TapeScreen assay (Agilent) and quantified by qPCR using the KAPA library quantification kit (Roche). Libraries were normalized to 2 nM and pooled for clustering on an Illumina cBot system using HiSeq PE Cluster Kit v4 reagents followed by sequencing on an Illumina HiSeq 2500 system with HiSeq SBS Kit v4 reagents with 100 bp paired-end reads.

### Mouse genome alignments and differential gene expression

Mapping to the mouse genome and differential expression analysis was conducted at AGRF under commercial contract using their in-house pipeline. The quality of the raw sequencing reads were assessed using FastQC and MultiQC. Adapters were trimmed using the TrimGalore (0.4.4) program and reads with a length <30 bp or quality <10 were removed. Filtered reads were aligned to the *Mus musculus* reference genome (mm10; GTF file GRCm38.6 – annotation release 105) using the STAR aligner (v2.5.3a) with default parameters plus a parameter to restrict multi-mapping reads (‘— outFilterMultimapNmax 2’). Counts per gene were summarized using the featureCounts (v1.4.6-p5) utility in Subread. A counts matrix was generated from the collective samples using in-house scripts and input to R (3.5.0) for differential expression analysis. Differential expression analysis was undertaken using EdgeR (3.22.3) with default settings and no filters, given the importance of key genes with low transcript abundance (Wilson *et al*., 2017) and the small percentage of cells infected by SCV-ZIKA/CHIK in the quadriceps muscles (with whole quadriceps muscles harvested for RNA-Seq). Counts were converted to relative counts (CPM) and normalized using the TMM method and modelled using the likelihood ratio test, glmLRT().

### Read alignments to the mouse and viral genomes

Sequencing reads were assessed using FastQC (Simons, 2010) (v0.11.8) and trimmed using Cutadapt (Martin, 2011) (v2.3) to remove adapter sequences and low-quality bases. Trimmed reads were aligned using STAR (Dobin *et al*., 2012) (v2.7.1a) to a combined reference that included the GRCm38 primary assembly and the GENCODE M23 gene model (Harrow *et al*., 2012), VACV Copenhagen (M35027.1; 191737 bp), ZIKV (KU321639.1; 10676 bp), and CHIKV (AM258992.1; 11601 bp). Quality control metrics were computed using RNA-SeQC (DeLuca *et al*., 2012) (v1.1.8) and RSeQC (Wang *et al*., 2012) (v3.0.0). SAMtools (Li *et al*., 2009a) (v1.9) was used to obtain alignments to the coding sequences of mature peptide features of VACV, CHIKV and ZIKV.

### Histology and immunohistochemistry

H&E staining was undertaken as described previously (Prow *et al*., 2019). IHC for neutrophils was undertaken as described (Prow *et al*., 2019) using Ly6G primary antibody (Abcam Anti-Mouse Neutrophil antibody Clone: NIMP-R14 cat. No. ab2557, Cambridge, UK) and Ly6 secondary antibody (Biocare Medical Rat on Mouse HRP Polymer cat. No. RT517L, Concord, CA USA). Apoptag staining used the Millipore Apoptag Peroxidase In Situ Apoptosis Detection kit (cat. No. S7100 Temecula, CA, USA). IHC for CHIKV capsid (monoclonal antibody 5.5G9 (Goh *et al*., 2015)) and ZIKA envelope (monoclonal antibody 4G2 (Hobson-Peters *et al*., 2019), was undertaken as described using NovaRed secondary antibody (Vector Laboratories ImmPACT NovaRed Peroxidase Substrate Kit cat. No. SK-4805 Burlingame, CA, USA). Slides were digitally scanned using Aperio AT Turbo (Leica Biosystems).

### GSEAs and Cytoscape

Gene Set Enrichment Analysis (GSEA) (Subramanian *et al*., 2005) was performed on a desktop application (GSEA v4.0.3) and the GenePattern Public server (Reich *et al*., 2006) using the “GSEAPreranked” module. Protein interaction networks of differentially expressed gene lists were visualized in Cytoscape (v3.7.2) (Shannon *et al*., 2003). Enrichment for biological processes, molecular functions, KEGG pathways and other gene ontology categories in DEG lists was elucidated using the STRING database (Szklarczyk *et al*., 2019).

### Statistics

Statistical analysis of experimental data was performed using IBM SPSS Statistics for Windows, Version 19.0 (IBM Corp., Armonk, New York, USA). The paired t-test was used when the difference in variances was <4, skewness was >2 and kurtosis was <2. Otherwise the non-parametric Wilcoxon Signed Rank tests was used.

## DATA AVAILABILITY

Datasets and analyses are available in the Supplementary Tables. All raw sequencing data (fastq files) was submitted to the Sequence Read Archive (SRA), BioProject accession: PRJNA610695.

## ACKNOWLEDGEMENTS

We thank the animal house staff and Histology and Imaging Services at QIMR B for their assistance. We thank Dr T Larcher (Institut National de Recherche Agronomique, France) for help interpreting histopathology. JEH was supported by a Research Training Program award for her PhD studies via the Faculty of Medicine at the University of Queensland (UQ). JEH also received funding from the Global Change Institute at UQ. NAP was awarded an Advance Queensland Research Fellowship by the Queensland Government, Australia, with co-funding from Sementis. AS is a Leadership Fellowship with the National Health and Medical Research Council of Australia, and is co-director of the AIDRC GVN Center of excellence.

## AUTHOR CONTRIBUTIONS

JEH, TD, ASl, SK, A-M P, LG performed the bioinformatics. TTL, NAP performed the experiments. PMH designed and provided access to the SCV-ZIKA/CHIK vaccine. PMH and KY generated supplementary annotation files. LL and JDH characterized and supplied the SCV vaccine. AS supervised the research, conceived the study and wrote the manuscript, with assistance from other authors.

## ADDITIONAL INFORMATION

Supplementary information accompanies the paper

## Competing interests

NAP, LL and JDH own Sementis shares. JDH is the current CSO of Sementis. AS was a consultant for Sementis. PMH was the previous CEO/CSO of Sementis. LL and NAP have had, and/or currently have, salary and/or project support from funds provided, whole or in part, via Sementis. Sementis had no role in the design and interpretation of the study, or in preparation of the manuscript.

## Supplementary figures

**S1 Fig.**
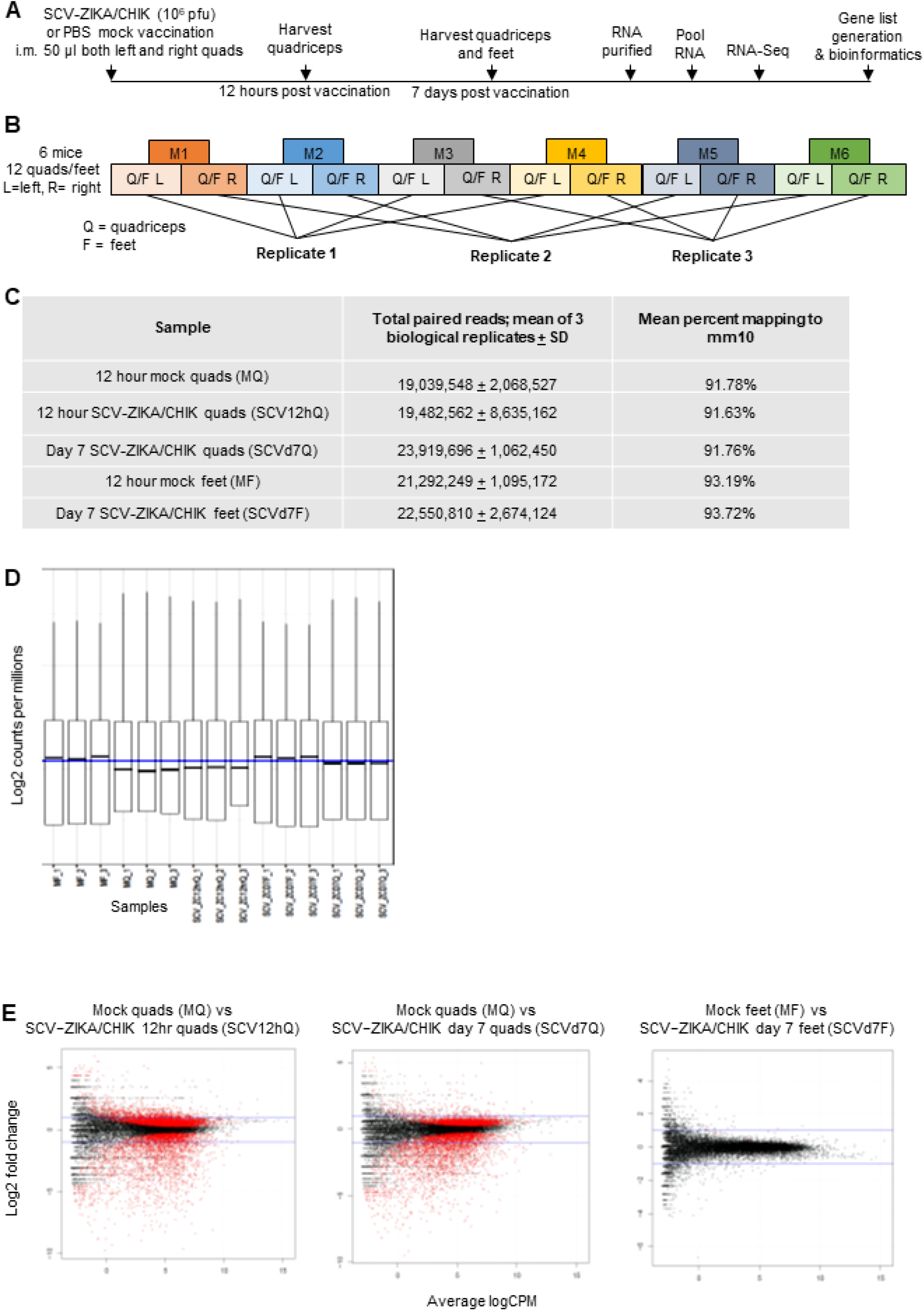
(A) Time line of experiment. (B) Pooling strategy for replicates. (C) Reads and percent of reads mapping to the mouse genome. (D) Boxplot of Log counts (normalized). Boxplots shows similar distributions of read counts amongst samples within and between groups. Boxes are 1^st^ & 3^rd^ quartile: whiskers range (no outliers). (E) Smear plots of the differentially expressed genes for the three comparisons. Red – FDR <0.05. Blue lines represent fold change of 2.

**S2 Fig.**
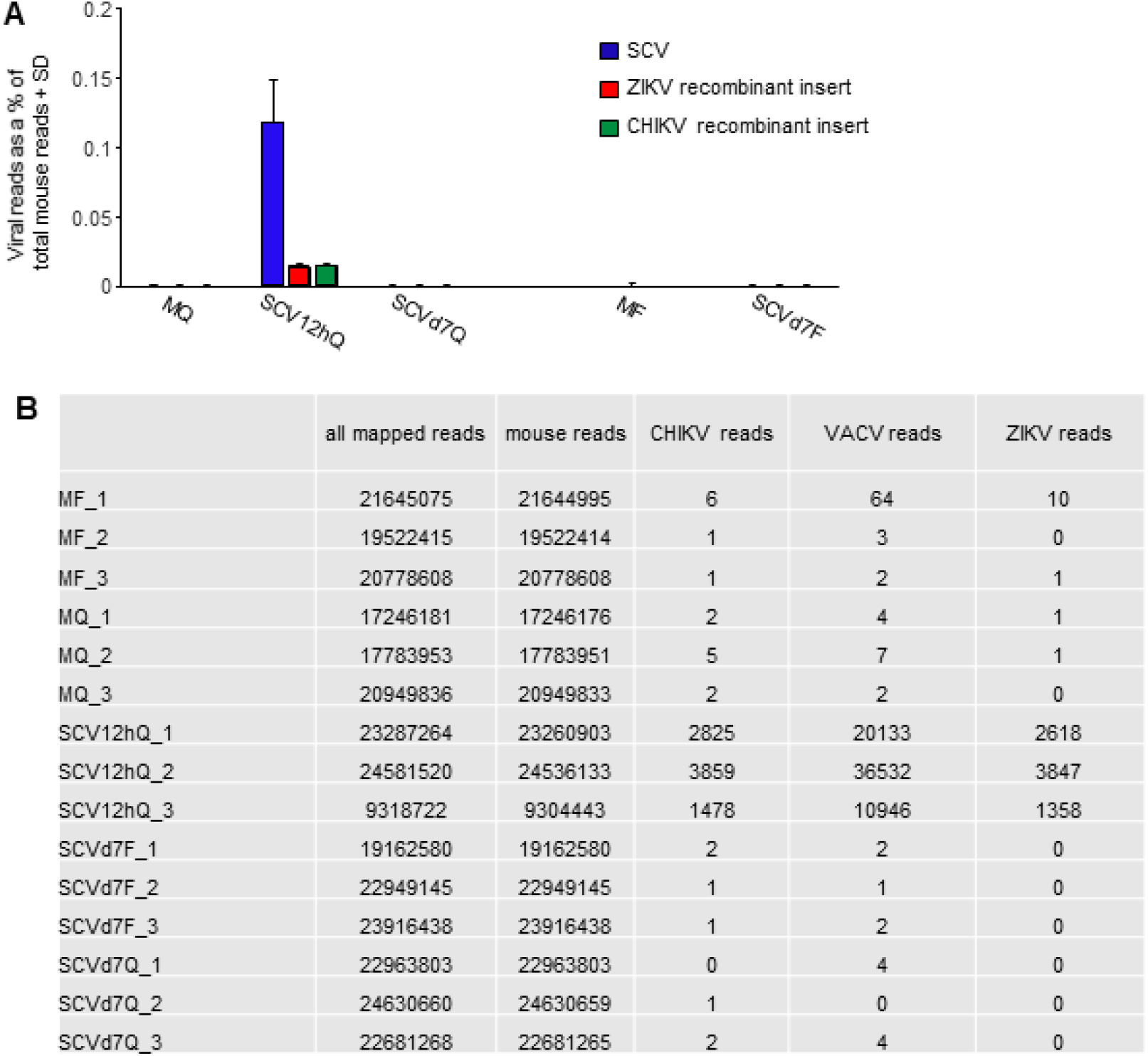
Bowtie2 alignments to viral genomes. Raw FASTQ files were assessed for quality using FastQC and MultiQC tools. Sequencing adapters were trimmed using Trimmomatic (0.36.6) ^1^ where reads with an average quality score over a 4 base sliding window of less than 20 were removed. Trimmed reads were aligned using Bowtie2 (v2.3.4.1) ^2^ to a combined reference that included the GRCm38 primary assembly and the GENCODE M23 gene model. Vaccinia virus Copenhagen (M35027.1), Zika virus strain ZikaSPH2015 (KU321639.1), and chikungunya virus (AM258992.1). Primary proper pair reads aligned to viral features, including CDS and mature peptide features, were counted using SAMtools (v1.9). **A** Bar graph of raw data shown in **B**. 1. Bolger AM, Lohse M, Usadel B. Trimmomatic: a flexible trimmer for Illumina sequence data. Bioinformatics. 2014:30(15)2114-20. 2. Langmead B, Salzberg SL. Fast gapped-read alignment with Bowtie 2. Nature Methods. 2012:9(4)357-9.

**S3 Fig.**
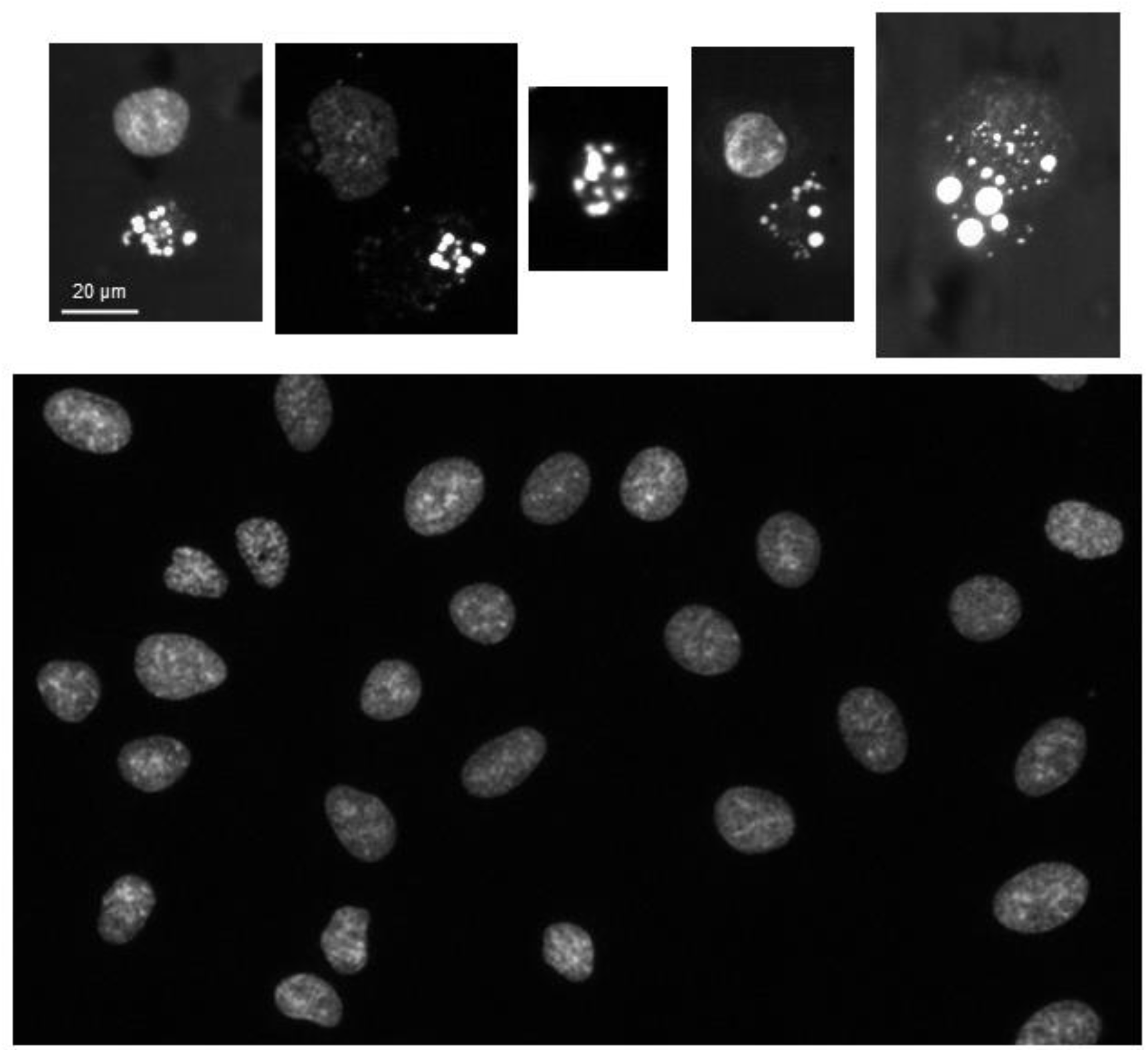
A549 cells infected with SCV-ZIKA/CHIK. 48-72 hours after SCV-ZIKA/CHIK infection, morphological features characteristic of apoptosis (condensation of chromatin) were clearly evident (top row) after staining with Hoechst 33342 (Linn *et al* J Virol Methods. 1995. 52(1-2):51-4). Bottom image shows uninfected controls.

## References

Albarnaz, J. D., Torres, A. A., & Smith, G. L. (2018). Modulating Vaccinia Virus Immunomodulators to Improve Immunological Memory. Viruses, 10(3), 101. doi:10.3390/v10030101

Alcam, A., Khanna, A., Paul, N. L., & Smith, G. L. (1999). Vaccinia virus strains Lister, USSR and Evans express soluble and cell-surface tumour necrosis factor receptors. J Gen Virol, 80 (Pt 4), 949–959. doi:10.1099/0022-1317-80-4-949

Alcami, A. (2020). Was smallpox a widespread mild disease? Science, 369(6502), 376–377. doi:10.1126/science.abd1214

Alcami, A., & Smith, G. L. (1992). A soluble receptor for interleukin-1 beta encoded by vaccinia virus: a novel mechanism of virus modulation of the host response to infection. Cell, 71(1), 153–167. doi:10.1016/0092-8674(92)90274-g

Alcamí, A., Symons, J. A., Collins, P. D., Williams, T. J., & Smith, G. L. (1998). Blockade of Chemokine Activity by a Soluble Chemokine Binding Protein from Vaccinia Virus. J Immunol, 160(2), 624–633.

Alharbi, N. K. (2019). Poxviral promoters for improving the immunogenicity of MVA delivered vaccines. Hum Vaccin Immunother, 15(1), 203–209. doi:10.1080/21645515.2018.1513439

Amsler, L., Malouli, D., & DeFilippis, V. (2013). The inflammasome as a target of modulation by DNA viruses. Future Virol, 8(4), 357–370. doi:10.2217/fvl.13.22

Bakshi, S., Taylor, J., Strickson, S., McCartney, T., & Cohen, P. (2017). Identification of TBK1 complexes required for the phosphorylation of IRF3 and the production of interferon beta. Biochem J, 474(7), 1163–1174. doi:10.1042/BCJ20160992

Barrera, J., Schutta, C., Pisano, M., Grubman, M. J., Brake, D. A., Miller, T., Kamicker, B. J., Olutunmbi, F., Ettyreddy, D., Brough, D. E., Butman, B. T., & Neilan, J. G. (2018). Use of ENABL(R) adjuvant to increase the potency of an adenovirus-vectored foot-and-mouth disease virus serotype A subunit vaccine. Vaccine, 36(8), 1078–1084. doi:10.1016/j.vaccine.2018.01.026

Bauer, S., Bathke, B., Lauterbach, H., Patzold, J., Kassub, R., Luber, C. A., Schlatter, B., Hamm, S., Chaplin, P., Suter, M., & Hochrein, H. (2010). A major role for TLR8 in the recognition of vaccinia viral DNA by murine pDC? Proc Natl Acad Sci U S A, 107(36), E139. doi:10.1073/pnas.1008626107

Beachkofsky, T. M., Carrizales, S. C., Bidinger, J. J., Hrncir, D. E., Whittemore, D. E., & Hivnor, C. M. (2010). Adverse events following smallpox vaccination with ACAM2000 in a military population. Arch Dermatol, 146(6), 656–661. doi:10.1001/archdermatol.2010.46

Bhalla, A., Zhao, N., Rivas, D. D., Ho, T., Perez de Llano, L., Mukherjee, M., & Nair, P. (2020). Exacerbations of severe asthma while on Anti-IL5 biologicals. J Investig Allergol Clin Immunol, 30(5). doi:10.18176/jiaci.0628

Bohnen, C., Wangorsch, A., Schulke, S., Nakajima-Adachi, H., Hachimura, S., Burggraf, M., Suzer, Y., Schwantes, A., Sutter, G., Waibler, Z., Reese, G., Toda, M., Scheurer, S., & Vieths, S. (2013). Vaccination with recombinant modified vaccinia virus Ankara prevents the onset of intestinal allergy in mice. Allergy, 68(8), 1021–1028. doi:10.1111/all.12192

Brandt, T. A., & Jacobs, B. L. (2001). Both carboxy- and amino-terminal domains of the vaccinia virus interferon resistance gene, E3L, are required for pathogenesis in a mouse model. J Virol, 75(2), 850–856. doi:10.1128/jvi.75.2.850-856.2001

Buchta, C. M., & Bishop, G. A. (2014). TRAF5 negatively regulates TLR signaling in B lymphocytes. J Immunol, 192(1), 145–150. doi:10.4049/jimmunol.1301901

Burns, J. M., Dairaghi, D. J., Deitz, M., Tsang, M., & Schall, T. J. (2002). Comprehensive mapping of poxvirus vCCI chemokine-binding protein. Expanded range of ligand interactions and unusual dissociation kinetics. J Biol Chem, 277(4), 2785–2789. doi:10.1074/jbc.M109884200

Carpenter, E. A., Ruby, J., & Ramshaw, I. A. (1994). IFN-gamma, TNF, and IL-6 production by vaccinia virus immune spleen cells. An in vitro study. J Immunol, 152(6), 2652–2659.

Carty, M., & Bowie, A. G. (2010). Recent insights into the role of Toll-like receptors in viral infection. Clin Exp Immunol, 161(3), 397–406. doi:10.1111/j.1365-2249.2010.04196.x

Chabot, V., Reverdiau, P., Iochmann, S., Rico, A., Senecal, D., Goupille, C., Sizaret, P. Y., & Sensebe, L. (2006). CCL5-enhanced human immature dendritic cell migration through the basement membrane in vitro depends on matrix metalloproteinase-9. J Leukoc Biol, 79(4), 767–778. doi:10.1189/jlb.0804464

Chan, C. Y., Chan, K. R., Chua, C. J., Nur Hazirah, S., Ghosh, S., Ooi, E. E., & Low, J. G. (2017). Early molecular correlates of adverse events following yellow fever vaccination. JCI Insight, 2(19), e96031. doi:10.1172/jci.insight.96031

Chea, L. S., Wyatt, L. S., Gangadhara, S., Moss, B., & Amara, R. R. (2019). Novel Modified Vaccinia Virus Ankara Vector Expressing Anti-apoptotic Gene B13R Delays Apoptosis and Enhances Humoral Responses. J Virol, 93(5), e01648–01618. doi:10.1128/JVI.01648-18

Clarke, E., Bashorun, A. O., Okoye, M., Umesi, A., Badjie Hydara, M., Adigweme, I., Dhere, R., Sethna, V., Kampmann, B., Goldblatt, D., Tate, A., Weiner, D. H., Flores, J., Alderson, M. R., & Lamola, S. (2020). Safety and immunogenicity of a novel 10-valent pneumococcal conjugate vaccine candidate in adults, toddlers, and infants in The Gambia-Results of a phase 1/2 randomized, double-blinded, controlled trial. Vaccine, 38(2), 399–410. doi:10.1016/j.vaccine.2019.08.072

Coden, M. E., & Berdnikovs, S. (2020). Eosinophils in wound healing and epithelial remodeling: Is coagulation a missing link? J Leukoc Biol, 108(1), 93–103. doi:10.1002/JLB.3MR0120-390R

Colamonici, O. R., Domanski, P., Sweitzer, S. M., Larner, A., & Buller, R. M. (1995). Vaccinia virus B18R gene encodes a type I interferon-binding protein that blocks interferon alpha transmembrane signaling. J Biol Chem, 270(27), 15974–15978. doi:10.1074/jbc.270.27.15974

Copland, A., Sparrow, A., Hart, P., Diogo, G. R., Paul, M., Azuma, M., & Reljic, R. (2019). Bacillus Calmette-Guerin Induces PD-L1 Expression on Antigen-Presenting Cells via Autocrine and Paracrine Interleukin-STAT3 Circuits. Sci Rep, 9(1), 3655. doi:10.1038/s41598-019-40145-0

Côté, A., Godbout, K., & Boulet, L.-P. (2020). The Management of Severe Asthma in 2020. Biochem Pharmacol, 179, 114112. doi:doi:10.1016/j.bcp.2020.114112.

Dai, P., Wang, W., Cao, H., Avogadri, F., Dai, L., Drexler, I., Joyce, J. A., Li, X. D., Chen, Z., Merghoub, T., Shuman, S., & Deng, L. (2014). Modified vaccinia virus Ankara triggers type I IFN production in murine conventional dendritic cells via a cGAS/STING-mediated cytosolic DNA-sensing pathway. PLoS Pathog, 10(4), e1003989. doi:10.1371/journal.ppat.1003989

Darling, A. R., Freyschmidt, E. J., Burton, O. T., Koleoglou, K. J., Oyoshi, M. K., & Oettgen, H. C. (2014). IL-10 suppresses IL-17-mediated dermal inflammation and reduces the systemic burden of Vaccinia virus in a mouse model of eczema vaccinatum. Clin Immunol, 150(2), 153–160. doi:10.1016/j.clim.2013.11.010

Davies, M. L., Sei, J. J., Siciliano, N. A., Xu, R. H., Roscoe, F., Sigal, L. J., Eisenlohr, L. C., & Norbury, C. C. (2014). MyD88-dependent immunity to a natural model of vaccinia virus infection does not involve Toll-like receptor 2. J Virol, 88(6), 3557–3567. doi:10.1128/JVI.02776-13

Day, A. J., & Milner, C. M. (2019). TSG-6: A multifunctional protein with anti-inflammatory and tissue-protective properties. Matrix Biol, 78-79, 60–83. doi:10.1016/j.matbio.2018.01.011

DeLuca, D. S., Levin, J. Z., Sivachenko, A., Fennell, T., Nazaire, M. D., Williams, C., Reich, M., Winckler, W., & Getz, G. (2012). RNA-SeQC: RNA-seq metrics for quality control and process optimization. Bioinformatics, 28(11), 1530–1532. doi:10.1093/bioinformatics/bts196

Dempsey, A., Keating, S. E., Carty, M., & Bowie, A. G. (2018). Poxviral protein E3-altered cytokine production reveals that DExD/H-box helicase 9 controls Toll-like receptor-stimulated immune responses. J Biol Chem, 293(39), 14989–15001. doi:10.1074/jbc.RA118.005089

Deng, L. (2017). The cytosolic DNA- and RNA-sensing pathways play important and non-redundant roles in host defense against vaccinia infection. J Immunol, 1987((1 Supplement) 148), 16.

DiPerna, G., Stack, J., Bowie, A. G., Boyd, A., Kotwal, G., Zhang, Z., Arvikar, S., Latz, E., Fitzgerald, K. A., & Marshall, W. L. (2004). Poxvirus protein N1L targets the I-kappaB kinase complex, inhibits signaling to NF-kappaB by the tumor necrosis factor superfamily of receptors, and inhibits NF-kappaB and IRF3 signaling by toll-like receptors. J Biol Chem, 279(35), 36570–36578. doi:10.1074/jbc.M400567200

Dobin, A., Davis, C. A., Schlesinger, F., Drenkow, J., Zaleski, C., Jha, S., Batut, P., Chaisson, M., & Gingeras, T. R. (2012). STAR: ultrafast universal RNA-seq aligner. Bioinformatics, 29(1), 15–21. doi:10.1093/bioinformatics/bts635

Earl, P. L., Hugin, A. W., & Moss, B. (1990). Removal of cryptic poxvirus transcription termination signals from the human immunodeficiency virus type 1 envelope gene enhances expression and immunogenicity of a recombinant vaccinia virus. J Virol, 64(5), 2448–2451.

Edelman, R., Tacket, C. O., Wasserman, S. S., Bodison, S. A., Perry, J. G., & Mangiafico, J. A. (2000). Phase II safety and immunogenicity study of live chikungunya virus vaccine TSI-GSD-218. Am J Trop Med Hyg, 62(6), 681–685. doi:10.4269/ajtmh.2000.62.681

Eldi, P., Cooper, T. H., Liu, L., Prow, N. A., Diener, K. R., Howley, P. M., Suhrbier, A., & Hayball, J. D. (2017). Production of a chikungunya vaccine using a CHO cell and attenuated viral-based platform technology. Mol Ther, 25(10), 2332–2344. doi:10.1016/j.ymthe.2017.06.017

Ferguson, B. J., Benfield, C. T. O., Ren, H., Lee, V. H., Frazer, G. L., Strnadova, P., Sumner, R. P., & Smith, G. L. (2013). Vaccinia virus protein N2 is a nuclear IRF3 inhibitor that promotes virulence. J Gen Virol, 94(Pt 9), 2070–2081. doi:10.1099/vir.0.054114-0

Flechsig, C., Suezer, Y., Kapp, M., Tan, S. M., Loffler, J., Sutter, G., Einsele, H., & Grigoleit, G. U. (2011). Uptake of antigens from modified vaccinia Ankara virus-infected leukocytes enhances the immunostimulatory capacity of dendritic cells. Cytotherapy, 13(6), 739–752. doi:10.3109/14653249.2010.549123

Forbester, J. L., Clement, M., Wellington, D., Yeung, A., Dimonte, S., Marsden, M., Chapman, L., Coomber, E. L., Tolley, C., Lees, E., Hale, C., Clare, S., Udalova, I., Dong, T., Dougan, G., & Humphreys, I. R. (2020). IRF5 Promotes Influenza Virus-Induced Inflammatory Responses in Human Induced Pluripotent Stem Cell-Derived Myeloid Cells and Murine Models. J Virol, 94(9), e00121–00120. doi:10.1128/JVI.00121-20

Forero, A., Ozarkar, S., Li, H., Lee, C. H., Hemann, E. A., Nadjsombati, M. S., Hendricks, M. R., So, L., Green, R., Roy, C. N., Sarkar, S. N., von Moltke, J., Anderson, S. K., Gale, M., Jr., & Savan, R. (2019). Differential Activation of the Transcription Factor IRF1 Underlies the Distinct Immune Responses Elicited by Type I and Type III Interferons. Immunity, 51(3), 451–464 e456. doi:10.1016/j.immuni.2019.07.007

Frey, S. E., Couch, R. B., Tacket, C. O., Treanor, J. J., Wolff, M., Newman, F. K., Atmar, R. L., Edelman, R., Nolan, C. M., Belshe, R. B., National Institute of, A., & Infectious Diseases Smallpox Vaccine Study, G. (2002). Clinical responses to undiluted and diluted smallpox vaccine. N Engl J Med, 346(17), 1265–1274. doi:10.1056/NEJMoa020534

Fulginiti, V. A., Papier, A., Lane, J. M., Neff, J. M., & Henderson, D. A. (2003). Smallpox vaccination: a review, part II. Adverse events. Clin Infect Dis, 37(2), 251–271. doi:10.1086/375825

Fuller, D. H., & Berglund, P. (2020). Amplifying RNA Vaccine Development. N Engl J Med, 382(25), 2469–2471. doi:10.1056/NEJMcibr2009737

García-Arriaza, J., Arnáez, P., Gómez, C. E., Sorzano, C. Ó. S., & Esteban, M. (2013). Improving Adaptive and Memory Immune Responses of an HIV/AIDS Vaccine Candidate MVA-B by Deletion of Vaccinia Virus Genes (C6L and K7R) Blocking Interferon Signaling Pathways. PLoS One, 8(6), e66894–e66894. doi:10.1371/journal.pone.0066894

Garcia-Arriaza, J., Gomez, C. E., Sorzano, C. O., & Esteban, M. (2014). Deletion of the vaccinia virus N2L gene encoding an inhibitor of IRF3 improves the immunogenicity of modified vaccinia virus Ankara expressing HIV-1 antigens. J Virol, 88(6), 3392–3410. doi:10.1128/JVI.02723-13

Gargett, T., Grubor-Bauk, B., Garrod, T. J., Yu, W., Miller, D., Major, L., Wesselingh, S., Suhrbier, A., & Gowans, E. J. (2014). Induction of antigen-positive cell death by the expression of perforin, but not DTa, from a DNA vaccine enhances the immune response. Immunol Cell Biol, 92(4), 359–367. doi:10.1038/icb.2013.93

Gatti-Mays, M. E., Strauss, J., Donahue, R. N., Palena, C., Del Rivero, J., Redman, J. M., Madan, R. A., Marte, J. L., Cordes, L. M., Lamping, E., Orpia, A., Burmeister, A., Wagner, E., Pico Navarro, C., Heery, C. R., Schlom, J., & Gulley, J. L. (2019). A Phase I Dose-Escalation Trial of BN-CV301, a Recombinant Poxviral Vaccine Targeting MUC1 and CEA with Costimulatory Molecules. Clin Cancer Res, 25(16), 4933–4944. doi:10.1158/1078-0432.CCR-19-0183

Gaylo-Moynihan, A., Prizant, H., Popovic, M., Fernandes, N. R. J., Anderson, C. S., Chiou, K. K., Bell, H., Schrock, D. C., Schumacher, J., Capece, T., Walling, B. L., Topham, D. J., Miller, J., Smrcka, A. V., Kim, M., Hughson, A., & Fowell, D. J. (2019). Programming of Distinct Chemokine-Dependent and -Independent Search Strategies for Th1 and Th2 Cells Optimizes Function at Inflamed Sites. Immunity, 51(2), 298–309.e296. doi:10.1016/j.immuni.2019.06.026

Georgana, I., Sumner, R. P., Towers, G. J., & Maluquer de Motes, C. (2018). Virulent Poxviruses Inhibit DNA Sensing by Preventing STING Activation. J Virol, 92(10), e02145–02117. doi:10.1128/JVI.02145-17

Gerlic, M., Faustin, B., Postigo, A., Yu, E. C., Proell, M., Gombosuren, N., Krajewska, M., Flynn, R., Croft, M., Way, M., Satterthwait, A., Liddington, R. C., Salek-Ardakani, S., Matsuzawa, S., & Reed, J. C. (2013). Vaccinia virus F1L protein promotes virulence by inhibiting inflammasome activation. Proc Natl Acad Sci U S A, 110(19), 7808–7813. doi:10.1073/pnas.1215995110

Geurts, J., Joosten, L. A., Takahashi, N., Arntz, O. J., Gluck, A., Bennink, M. B., van den Berg, W. B., & van de Loo, F. A. (2009). Computational design and application of endogenous promoters for transcriptionally targeted gene therapy for rheumatoid arthritis. Mol Ther, 17(11), 1877–1887. doi:10.1038/mt.2009.182

Gholobova, D., Gerard, M., Terrie, L., Desender, L., Shansky, J., Vandenburgh, H., & Thorrez, L. (2019). Coculture Method to Obtain Endothelial Networks Within Human Tissue-Engineered Skeletal Muscle. Methods Mol Biol, 1889, 169–183. doi:10.1007/978-1-4939-8897-6_10

Goh, L. Y. H., Hobson-Peters, J., Prow, N. A., Gardner, J., Bielefeldt-Ohmann, H., Suhrbier, A., & Hall, R. A. (2015). Monoclonal antibodies specific for the capsid protein of chikungunya virus suitable for multiple applications. J Gen Virol, 96(Pt 3), 507–512. doi:10.1099/jgv.0.000002

Gomes, A. C., Mohsen, M. O., Mueller, J. E., Leoratti, F. M. S., Cabral-Miranda, G., & Bachmann, M. F. (2019). Early Transcriptional Signature in Dendritic Cells and the Induction of Protective T Cell Responses Upon Immunization With VLPs Containing TLR Ligands-A Role for CCL2. Front Immunol, 10, 1679. doi:10.3389/fimmu.2019.01679

Gonzalez-Dias, P., Lee, E. K., Sorgi, S., de Lima, D. S., Urbanski, A. H., Silveira, E. L., & Nakaya, H. I. (2020). Methods for predicting vaccine immunogenicity and reactogenicity. Hum Vaccin Immunother, 16(2), 269–276. doi:10.1080/21645515.2019.1697110

Gray, J. E., Chiappori, A., Williams, C. C., Tanvetyanon, T., Haura, E. B., Creelan, B. C., Kim, J., Boyle, T. A., Pinder-Schenck, M., Khalil, F., Altiok, S., Devane, R., Noyes, D., Mediavilla-Varela, M., Smilee, R., Hopewell, E. L., Kelley, L., & Antonia, S. J. (2018). A phase I/randomized phase II study of GM.CD40L vaccine in combination with CCL21 in patients with advanced lung adenocarcinoma. Cancer Immunol Immunother, 67(12), 1853–1862. doi:10.1007/s00262-018-2236-7

Harenberg, A., Guillaume, F., Ryan, E. J., Burdin, N., & Spada, F. (2008). Gene profiling analysis of ALVAC infected human monocyte derived dendritic cells. Vaccine, 26(39), 5004–5013. doi:10.1016/j.vaccine.2008.07.050

Harrington, L. E., Most Rv, R., Whitton, J. L., & Ahmed, R. (2002). Recombinant vaccinia virus-induced T-cell immunity: quantitation of the response to the virus vector and the foreign epitope. J Virol, 76(7), 3329–3337. doi:10.1128/jvi.76.7.3329-3337.2002

Harrow, J., Frankish, A., Gonzalez, J. M., Tapanari, E., Diekhans, M., Kokocinski, F., Aken, B. L., Barrell, D., Zadissa, A., Searle, S., Barnes, I., Bignell, A., Boychenko, V., Hunt, T., Kay, M., Mukherjee, G., Rajan, J., Despacio-Reyes, G., Saunders, G., Steward, C., Harte, R., Lin, M., Howald, C., Tanzer, A., Derrien, T., Chrast, J., Walters, N., Balasubramanian, S., Pei, B., Tress, M., Rodriguez, J. M., Ezkurdia, I., van Baren, J., Brent, M., Haussler, D., Kellis, M., Valencia, A., Reymond, A., Gerstein, M., Guigo, R., & Hubbard, T. J. (2012). GENCODE: the reference human genome annotation for The ENCODE Project. Genome Res, 22(9), 1760–1774. doi:10.1101/gr.135350.111

Hatesuer, B., Hoang, H. T., Riese, P., Trittel, S., Gerhauser, I., Elbahesh, H., Geffers, R., Wilk, E., & Schughart, K. (2017). Deletion of Irf3 and Irf7 Genes in Mice Results in Altered Interferon Pathway Activation and Granulocyte-Dominated Inflammatory Responses to Influenza A Infection. J Innate Immun, 9(2), 145–161. doi:10.1159/000450705

He, Y., Fisher, R., Chowdhury, S., Sultana, I., Pereira, C. P., Bray, M., & Reed, J. L. (2014). Vaccinia virus induces rapid necrosis in keratinocytes by a STAT3-dependent mechanism. PLoS One, 9(11), e113690. doi:10.1371/journal.pone.0113690

Heredia, J. E., Mukundan, L., Chen, F. M., Mueller, A. A., Deo, R. C., Locksley, R. M., Rando, T. A., & Chawla, A. (2013). Type 2 innate signals stimulate fibro/adipogenic progenitors to facilitate muscle regeneration. Cell, 153(2), 376–388. doi:10.1016/j.cell.2013.02.053

Herve, C., Laupeze, B., Del Giudice, G., Didierlaurent, A. M., & Tavares Da Silva, F. (2019). The how’s and what’s of vaccine reactogenicity. NPJ Vaccines, 4, 39. doi:10.1038/s41541-019-0132-6

Hobson-Peters, J., Harrison, J. J., Watterson, D., Hazlewood, J. E., Vet, L. J., Newton, N. D., Warrilow, D., Colmant, A. M. G., Taylor, C., Huang, B., Piyasena, T. B. H., Chow, W. K., Setoh, Y. X., Tang, B., Nakayama, E., Yan, K., Amarilla, A. A., Wheatley, S., Moore, P. R., Finger, M., Kurucz, N., Modhiran, N., Young, P. R., Khromykh, A. A., Bielefeldt-Ohmann, H., Suhrbier, A., & Hall, R. A. (2019). A recombinant platform for flavivirus vaccines and diagnostics using chimeras of a new insect-specific virus. Sci Transl Med, 11(522), eaax7888. doi:10.1126/scitranslmed.aax7888

Hsu, K. L., Tsuboi, K., Adibekian, A., Pugh, H., Masuda, K., & Cravatt, B. F. (2012). DAGLbeta inhibition perturbs a lipid network involved in macrophage inflammatory responses. Nat Chem Biol, 8(12), 999–1007. doi:10.1038/nchembio.1105

Hutchens, M., Luker, K. E., Sottile, P., Sonstein, J., Lukacs, N. W., Nunez, G., Curtis, J. L., & Luker, G. D. (2008a). TLR3 increases disease morbidity and mortality from vaccinia infection. J Immunol, 180(1), 483–491. doi:10.4049/jimmunol.180.1.483

Hutchens, M. A., Luker, K. E., Sonstein, J., Núñez, G., Curtis, J. L., & Luker, G. D. (2008b). Protective Effect of Toll-like Receptor 4 in Pulmonary Vaccinia Infection. PLOS Pathogens, 4(9), e1000153. doi:10.1371/journal.ppat.1000153

IMVANEX: European Medicines Agency. Product Information. Updated 24/6/2020. https://www.ema.europa.eu/en/documents/product-information/imvanex-epar-product-information_en.pdf.

Ink, B. S., Gilbert, C. S., & Evan, G. I. (1995). Delay of vaccinia virus-induced apoptosis in nonpermissive Chinese hamster ovary cells by the cowpox virus CHOhr and adenovirus E1B 19K genes. J Virol, 69(2), 661–668. doi:doi:10.1128/JVI.69.2.661-668.1995

Izzi, V., Buler, M., Masuelli, L., Giganti, M. G., Modesti, A., & Bei, R. (2014). Poxvirus-based vaccines for cancer immunotherapy: new insights from combined cytokines/co-stimulatory molecules delivery and “uncommon” strains. Anticancer Agents Med Chem, 14(2), 183–189. doi:10.2174/18715206113136660376

Jackson, S. S., Ilyinskii, P., Philippon, V., Gritz, L., Yafal, A. G., Zinnack, K., Beaudry, K. R., Manson, K. H., Lifton, M. A., Kuroda, M. J., Letvin, N. L., Mazzara, G. P., & Panicali, D. L. (2005). Role of genes that modulate host immune responses in the immunogenicity and pathogenicity of vaccinia virus. J Virol, 79(10), 6554–6559. doi:10.1128/JVI.79.10.6554-6559.2005

James, K. R., Soon, M. S. F., Sebina, I., Fernandez-Ruiz, D., Davey, G., Liligeto, U. N., Nair, A. S., Fogg, L. G., Edwards, C. L., Best, S. E., Lansink, L. I. M., Schroder, K., Wilson, J. A. C., Austin, R., Suhrbier, A., Lane, S. W., Hill, G. R., Engwerda, C. R., Heath, W. R., & Haque, A. (2018). IFN Regulatory Factor 3 Balances Th1 and T Follicular Helper Immunity during Nonlethal Blood-Stage Plasmodium Infection. J Immunol, 200(4), 1443–1456. doi:10.4049/jimmunol.1700782

Jefferies, C. A. (2019). Regulating IRFs in IFN Driven Disease. Front Immunol, 10, 325. doi:10.3389/fimmu.2019.00325

Jiang, M., Chen, P., Wang, L., Li, W., Chen, B., Liu, Y., Wang, H., Zhao, S., Ye, L., He, Y., & Zhou, C. (2020). cGAS-STING, an important pathway in cancer immunotherapy. J Hematol Oncol, 13(1), 81. doi:10.1186/s13045-020-00916-z

Joachim, A., Ahmed, M. I. M., Pollakis, G., Rogers, L., Hoffmann, V. S., Munseri, P., Aboud, S., Lyamuya, E. F., Bakari, M., Robb, M. L., Wahren, B., Sandstrom, E., Nilsson, C., Biberfeld, G., Geldmacher, C., & Held, K. (2020). Induction of Identical IgG HIV-1 Envelope Epitope Recognition Patterns After Initial HIVIS-DNA/MVA-CMDR Immunization and a Late MVA-CMDR Boost. Front Immunol, 11, 719. doi:10.3389/fimmu.2020.00719

Katsafanas, G. C., & Moss, B. (2007). Colocalization of transcription and translation within cytoplasmic poxvirus factories coordinates viral expression and subjugates host functions. Cell Host Microbe, 2(4), 221–228. doi:10.1016/j.chom.2007.08.005

Kawasaki, T., & Kawai, T. (2014). Toll-like receptor signaling pathways. Front Immunol, 5, 461. doi:10.3389/fimmu.2014.00461

Kelley, N., Jeltema, D., Duan, Y., & He, Y. (2019). The NLRP3 Inflammasome: An Overview of Mechanisms of Activation and Regulation. Int J Mol Sci, 20(13), 3328. doi:10.3390/ijms20133328

Kieser, Q., Noyce, R. S., Shenouda, M., Lin, Y. J., & Evans, D. H. (2020). Cytoplasmic factories, virus assembly, and DNA replication kinetics collectively constrain the formation of poxvirus recombinants. PLoS One, 15(1), e0228028. doi:10.1371/journal.pone.0228028

Kluczyk, A., Siemion, I. Z., Szewczuk, Z., & Wieczorek, Z. (2002). The immunosuppressive activity of peptide fragments of vaccinia virus C10L protein and a hypothesis on the role of this protein in the viral invasion. Peptides, 23(5), 823–834. doi:10.1016/s0196-9781(02)00006-2

Kobayashi, K., Hernandez, L. D., Galan, J. E., Janeway, C. A., Jr., Medzhitov, R., & Flavell, R. A. (2002). IRAK-M is a negative regulator of Toll-like receptor signaling. Cell, 110(2), 191–202. doi:10.1016/s0092-8674(02)00827-9

Koch, T., Dahlke, C., Fathi, A., Kupke, A., Krähling, V., Okba, N. M. A., Halwe, S., Rohde, C., Eickmann, M., Volz, A., Hesterkamp, T., Jambrecina, A., Borregaard, S., Ly, M. L., Zinser, M. E., Bartels, E., Poetsch, J. S. H., Neumann, R., Fux, R., Schmiedel, S., Lohse, A. W., Haagmans, B. L., Sutter, G., Becker, S., & Addo, M. M. (2020). Safety and immunogenicity of a modified vaccinia virus Ankara vector vaccine candidate for Middle East respiratory syndrome: an open-label, phase 1 trial. Lancet Infect Dis, 20(7), 827–838. doi:10.1016/S1473-3099(20)30248-6

Kohonen-Corish, M. R., King, N. J., Woodhams, C. E., & Ramshaw, I. A. (1990). Immunodeficient mice recover from infection with vaccinia virus expressing interferon-gamma. Eur J Immunol, 20(1), 157–161. doi:10.1002/eji.1830200123

Laher, F., Moodie, Z., Cohen, K. W., Grunenberg, N., Bekker, L. G., Allen, M., Frahm, N., Yates, N. L., Morris, L., Malahleha, M., Mngadi, K., Daniels, B., Innes, C., Saunders, K., Grant, S., Yu, C., Gilbert, P. B., Phogat, S., DiazGranados, C. A., Koutsoukos, M., Van Der Meeren, O., Bentley, C., Mkhize, N. N., Pensiero, M. N., Mehra, V. L., Kublin, J. G., Corey, L., Montefiori, D. C., Gray, G. E., McElrath, M. J., & Tomaras, G. D. (2020). Safety and immune responses after a 12-month booster in healthy HIV-uninfected adults in HVTN 100 in South Africa: A randomized double-blind placebo-controlled trial of ALVAC-HIV (vCP2438) and bivalent subtype C gp120/MF59 vaccines. PLoS Med, 17(2), e1003038. doi:10.1371/journal.pmed.1003038

Langland, J. O., Kash, J. C., Carter, V., Thomas, M. J., Katze, M. G., & Jacobs, B. L. (2006). Suppression of proinflammatory signal transduction and gene expression by the dual nucleic acid binding domains of the vaccinia virus E3L proteins. J Virol, 80(20), 10083–10095. doi:10.1128/JVI.00607-06

Lee, C. S., Lee, K. H., Jung, M. H., & Lee, H. B. (2008). Rate of influenza vaccination and its adverse reactions seen in health care personnel in a single tertiary hospital in Korea. Jpn J Infect Dis, 61(6), 457–460.

Legrand, F. A., Verardi, P. H., Jones, L. A., Chan, K. S., Peng, Y., & Yilma, T. D. (2004). Induction of potent humoral and cell-mediated immune responses by attenuated vaccinia virus vectors with deleted serpin genes. J Virol, 78(6), 2770–2779. doi:10.1128/jvi.78.6.2770-2779.2004

Lehmann, M. H., Kastenmuller, W., Kandemir, J. D., Brandt, F., Suezer, Y., & Sutter, G. (2009). Modified vaccinia virus ankara triggers chemotaxis of monocytes and early respiratory immigration of leukocytes by induction of CCL2 expression. J Virol, 83(6), 2540–2552. doi:10.1128/JVI.01884-08

Lehmann, M. H., Torres-Domínguez, L. E., Price, P. J., Brandmüller, C., Kirschning, C. J., & Sutter, G. (2016). CCL2 expression is mediated by type I IFN receptor and recruits NK and T cells to the lung during MVA infection. J Leukoc Biol, 99(6), 1057–1064. doi:10.1189/jlb.4MA0815-376RR

Lei, B. U. W., & Prow, T. W. (2019). A review of microsampling techniques and their social impact. Biomed Microdevices, 21(4), 81. doi:10.1007/s10544-019-0412-y

Lentscher, A. J., McCarthy, M. K., May, N. A., Davenport, B. J., Montgomery, S. A., Raghunathan, K., McAllister, N., Silva, L. A., Morrison, T. E., & Dermody, T. S. (2020). Chikungunya virus replication in skeletal muscle cells is required for disease development. J Clin Invest, 130(3), 1466–1478. doi:10.1172/JCI129893

Li, H., Handsaker, B., Wysoker, A., Fennell, T., Ruan, J., Homer, N., Marth, G., Abecasis, G., Durbin, R., & Genome Project Data Processing, S. (2009a). The Sequence Alignment/Map format and SAMtools. Bioinformatics, 25(16), 2078–2079. doi:10.1093/bioinformatics/btp352

Li, S., Rouphael, N., Duraisingham, S., Romero-Steiner, S., Presnell, S., Davis, C., Schmidt, D. S., Johnson, S. E., Milton, A., Rajam, G., Kasturi, S., Carlone, G. M., Quinn, C., Chaussabel, D., Palucka, A. K., Mulligan, M. J., Ahmed, R., Stephens, D. S., Nakaya, H. I., & Pulendran, B. (2014). Molecular signatures of antibody responses derived from a systems biology study of five human vaccines. Nat Immunol, 15(2), 195–204. doi:10.1038/ni.2789

Li, Y., Cao, C., Jia, W., Yu, L., Mo, M., Wang, Q., Huang, Y., Lim, J. M., Ishihara, M., Wells, L., Azadi, P., Robinson, H., He, Y. W., Zhang, L., & Mariuzza, R. A. (2009b). Structure of the F-spondin domain of mindin, an integrin ligand and pattern recognition molecule. EMBO J, 28(3), 286–297. doi:10.1038/emboj.2008.288

Li, Y., Meyer, H., Zhao, H., & Damon, I. K. (2010). GC content-based pan-pox universal PCR assays for poxvirus detection. J Clin Microbiol, 48(1), 268–276. doi:10.1128/JCM.01697-09

Liu, F., Niu, Q., Fan, X., Liu, C., Zhang, J., Wei, Z., Hou, W., Kanneganti, T. D., Robb, M. L., Kim, J. H., Michael, N. L., Sun, J., Soong, L., & Hu, H. (2017). Priming and Activation of Inflammasome by Canarypox Virus Vector ALVAC via the cGAS/IFI16-STING-Type I IFN Pathway and AIM2 Sensor. J Immunol, 199(9), 3293–3305. doi:10.4049/jimmunol.1700698

Liu, S. W., Katsafanas, G. C., Liu, R., Wyatt, L. S., & Moss, B. (2015). Poxvirus decapping enzymes enhance virulence by preventing the accumulation of dsRNA and the induction of innate antiviral responses. Cell Host Microbe, 17(3), 320–331. doi:10.1016/j.chom.2015.02.002

Lloyd, C. M., & Snelgrove, R. J. (2018). Type 2 immunity: Expanding our view. Sci Immunol, 3(25), eaat1604. doi:10.1126/sciimmunol.aat1604

Lopez-Gil, E., Lorenzo, G., Hevia, E., Borrego, B., Eiden, M., Groschup, M., Gilbert, S. C., & Brun, A. (2013). A single immunization with MVA expressing GnGc glycoproteins promotes epitope-specific CD8+-T cell activation and protects immune-competent mice against a lethal RVFV infection. PLoS Negl Trop Dis, 7(7), e2309. doi:10.1371/journal.pntd.0002309

Lopez, M. J., Seyed-Razavi, Y., Jamali, A., Harris, D. L., & Hamrah, P. (2018). The Chemokine Receptor CXCR4 Mediates Recruitment of CD11c+ Conventional Dendritic Cells Into the Inflamed Murine Cornea. Invest Ophthalmol Vis Sci, 59(13), 5671–5681. doi:10.1167/iovs.18-25084

Lousberg, E. L., Diener, K. R., Fraser, C. K., Phipps, S., Foster, P. S., Chen, W., Uematsu, S., Akira, S., Robertson, S. A., Brown, M. P., & Hayball, J. D. (2011). Antigen-specific T-cell responses to a recombinant fowlpox virus are dependent on MyD88 and interleukin-18 and independent of Toll-like receptor 7 (TLR7)- and TLR9-mediated innate immune recognition. J Virol, 85(7), 3385–3396. doi:10.1128/JVI.02000-10

Luo, J., Liu, X. P., Xiong, F. F., Gao, F. X., Yi, Y. L., Zhang, M., Chen, Z., & Tan, W. S. (2019). Enhancing Immune Response and Heterosubtypic Protection Ability of Inactivated H7N9 Vaccine by Using STING Agonist as a Mucosal Adjuvant. Front Immunol, 10, 2274. doi:10.3389/fimmu.2019.02274

Lysakova-Devine, T., Keogh, B., Harrington, B., Nagpal, K., Halle, A., Golenbock, D. T., Monie, T., & Bowie, A. G. (2010). Viral inhibitory peptide of TLR4, a peptide derived from vaccinia protein A46, specifically inhibits TLR4 by directly targeting MyD88 adaptor-like and TRIF-related adaptor molecule. J Immunol, 185(7), 4261–4271. doi:10.4049/jimmunol.1002013

Magnusson, S. E., Altenburg, A. F., Bengtsson, K. L., Bosman, F., de Vries, R. D., Rimmelzwaan, G. F., & Stertman, L. (2018). Matrix-M adjuvant enhances immunogenicity of both protein- and modified vaccinia virus Ankara-based influenza vaccines in mice. Immunol Res, 66(2), 224–233. doi:10.1007/s12026-018-8991-x

Maluquer de Motes, C., Cooray, S., Ren, H., Almeida, G. M. F., McGourty, K., Bahar, M. W., Stuart, D. I., Grimes, J. M., Graham, S. C., & Smith, G. L. (2011). Inhibition of Apoptosis and NF-κB Activation by Vaccinia Protein N1 Occur via Distinct Binding Surfaces and Make Different Contributions to Virulence. PLOS Pathogens, 7(12), e1002430. doi:10.1371/journal.ppat.1002430

Mann, B. A., Huang, J. H., Li, P., Chang, H.-C., Slee, R. B., O’Sullivan, A., Anita, M., Yeh, N., Klemsz, M. J., Brutkiewicz, R. R., Blum, J. S., & Kaplan, M. H. (2008). Vaccinia virus blocks Stat1-dependent and Stat1-independent gene expression induced by type I and type II interferons. J Interferon Cytokine Res, 28(6), 367–380. doi:10.1089/jir.2007.0113

Marin, M. Q., Perez, P., Gomez, C. E., Sorzano, C. O. S., Esteban, M., & Garcia-Arriaza, J. (2018). Removal of the C6 Vaccinia Virus Interferon-beta Inhibitor in the Hepatitis C Vaccine Candidate MVA-HCV Elicited in Mice High Immunogenicity in Spite of Reduced Host Gene Expression. Viruses, 10(8). doi:10.3390/v10080414

Martin-Sanchez, F., Diamond, C., Zeitler, M., Gomez, A. I., Baroja-Mazo, A., Bagnall, J., Spiller, D., White, M., Daniels, M. J., Mortellaro, A., Penalver, M., Paszek, P., Steringer, J. P., Nickel, W., Brough, D., & Pelegrin, P. (2016). Inflammasome-dependent IL-1beta release depends upon membrane permeabilisation. Cell Death Differ, 23(7), 1219–1231. doi:10.1038/cdd.2015.176

Martin, M. (2011). Cutadapt removes adapter sequences from high-throughput sequencing reads. EMBnet.journal, 17(1). doi:https://doi.org/10.14806/ej.17.1.200

Martinez, J., Huang, X., & Yang, Y. (2010). Direct TLR2 signaling is critical for NK cell activation and function in response to vaccinia viral infection. PLoS Pathog, 6(3), e1000811. doi:10.1371/journal.ppat.1000811

Martins, K. A., Bavari, S., & Salazar, A. M. (2015). Vaccine adjuvant uses of poly-IC and derivatives. Expert Rev Vaccines, 14(3), 447–459. doi:10.1586/14760584.2015.966085

Matchett, W. E., Malewana, G. B. R., Mudrick, H., Medlyn, M. J., & Barry, M. A. (2020). Genetic Adjuvants in Replicating Single-Cycle Adenovirus Vectors Amplify Systemic and Mucosal Immune Responses against HIV-1 Envelope. Vaccines (Basel), 8(1), 64. doi:10.3390/vaccines8010064

Matthijs, A. M. F., Auray, G., Jakob, V., Garcia-Nicolas, O., Braun, R. O., Keller, I., Bruggman, R., Devriendt, B., Boyen, F., Guzman, C. A., Michiels, A., Haesebrouck, F., Collin, N., Barnier-Quer, C., Maes, D., & Summerfield, A. (2019). Systems Immunology Characterization of Novel Vaccine Formulations for Mycoplasma hyopneumoniae Bacterins. Front Immunol, 10, 1087. doi:10.3389/fimmu.2019.01087

McKay, P. F., Cizmeci, D., Aldon, Y., Maertzdorf, J., Weiner, J., Kaufmann, S. H., Lewis, D. J., van den Berg, R. A., Del Giudice, G., & Shattock, R. J. (2019). Identification of potential biomarkers of vaccine inflammation in mice. Elife, 8, e46149. doi:10.7554/eLife.46149

Meixiong, J., & Dong, X. (2017). Mas-Related G Protein-Coupled Receptors and the Biology of Itch Sensation. Annu Rev Genet, 51, 103–121. doi:10.1146/annurev-genet-120116-024723

Michalska, A., Blaszczyk, K., Wesoly, J., & Bluyssen, H. A. R. (2018). A Positive Feedback Amplifier Circuit That Regulates Interferon (IFN)-Stimulated Gene Expression and Controls Type I and Type II IFN Responses. Front Immunol, 9, 1135. doi:10.3389/fimmu.2018.01135

Michlmayr, D., Pak, T. R., Rahman, A. H., Amir, E. D., Kim, E. Y., Kim-Schulze, S., Suprun, M., Stewart, M. G., Thomas, G. P., Balmaseda, A., Wang, L., Zhu, J., Suarez-Farinas, M., Wolinsky, S. M., Kasarskis, A., & Harris, E. (2018). Comprehensive innate immune profiling of chikungunya virus infection in pediatric cases. Mol Syst Biol, 14(8), e7862. doi:10.15252/msb.20177862

Morgan, J. E., Prola, A., Mariot, V., Pini, V., Meng, J., Hourde, C., Dumonceaux, J., Conti, F., Relaix, F., Authier, F. J., Tiret, L., Muntoni, F., & Bencze, M. (2018). Necroptosis mediates myofibre death in dystrophin-deficient mice. Nat Commun, 9(1), 3655. doi:10.1038/s41467-018-06057-9

Moss, B., Ahn, B. Y., Amegadzie, B., Gershon, P. D., & Keck, J. G. (1991). Cytoplasmic transcription system encoded by vaccinia virus. J Biol Chem, 266(3), 1355–1358.

Muhlemann, B., Vinner, L., Margaryan, A., Wilhelmson, H., de la Fuente Castro, C., Allentoft, M. E., de Barros Damgaard, P., Hansen, A. J., Holtsmark Nielsen, S., Strand, L. M., Bill, J., Buzhilova, A., Pushkina, T., Falys, C., Khartanovich, V., Moiseyev, V., Jorkov, M. L. S., Ostergaard Sorensen, P., Magnusson, Y., Gustin, I., Schroeder, H., Sutter, G., Smith, G. L., Drosten, C., Fouchier, R. A. M., Smith, D. J., Willerslev, E., Jones, T. C., & Sikora, M. (2020). Diverse variola virus (smallpox) strains were widespread in northern Europe in the Viking Age. Science, 369(6502), eaaw8977. doi:10.1126/science.aaw8977

Munoz-Wolf, N., & Lavelle, E. C. (2018). A Guide to IL-1 family cytokines in adjuvanticity. FEBS J, 285(13), 2377–2401. doi:10.1111/febs.14467

Myskiw, C., Arsenio, J., Booy, E. P., Hammett, C., Deschambault, Y., Gibson, S. B., & Cao, J. (2011). RNA species generated in vaccinia virus infected cells activate cell type-specific MDA5 or RIG-I dependent interferon gene transcription and PKR dependent apoptosis. Virology, 413(2), 183–193. doi:10.1016/j.virol.2011.01.034

Nagata, L. P., Irwin, C. R., Hu, W. G., & Evans, D. H. (2018). Vaccinia-based vaccines to biothreat and emerging viruses. Biotechnol Genet Eng Rev, 34(1), 107–121. doi:10.1080/02648725.2018.1471643

Nair, S. R., Abraham, R., Sundaram, S., & Sreekumar, E. (2017). Interferon regulated gene (IRG) expression-signature in a mouse model of chikungunya virus neurovirulence. J Neurovirol, 23(6), 886–902. doi:10.1007/s13365-017-0583-3

Natrajan, M. S., Rouphael, N., Lai, L., Kazmin, D., Jensen, T. L., Weiss, D. S., Ibegbu, C., Sztein, M. B., Hooper, W. F., Hill, H., Anderson, E. J., Johnson, R., Sanz, P., Pulendran, B., Goll, J. B., & Mulligan, M. J. (2019). Systems Vaccinology for a Live Attenuated Tularemia Vaccine Reveals Unique Transcriptional Signatures That Predict Humoral and Cellular Immune Responses. Vaccines (Basel), 8(1), 4. doi:10.3390/vaccines8010004

Negishi, H., Ohba, Y., Yanai, H., Takaoka, A., Honma, K., Yui, K., Matsuyama, T., Taniguchi, T., & Honda, K. (2005). Negative regulation of Toll-like-receptor signaling by IRF-4. Proc Natl Acad Sci U S A, 102(44), 15989–15994. doi:10.1073/pnas.0508327102

Neidel, S., Ren, H., Abreu Torres, A., & Smith, G. (2019). NF-κB activation is a turn on for vaccinia virus phosphoprotein A49 to turn off NF-κB activation. Proc Natl Acad Sci U S A, 116(12), 5699–5704. doi:10.1073/pnas.1813504116

Ng, C. S., Kato, H., & Fujita, T. (2019a). Fueling Type I Interferonopathies: Regulation and Function of Type I Interferon Antiviral Responses. J Interferon Cytokine Res, 39(7), 383–392. doi:10.1089/jir.2019.0037

Ng, H. I., Tuong, Z. K., Fernando, G. J. P., Depelsenaire, A. C. I., Meliga, S. C., Frazer, I. H., & Kendall, M. A. F. (2019b). Microprojection arrays applied to skin generate mechanical stress, induce an inflammatory transcriptome and cell death, and improve vaccine-induced immune responses. NPJ Vaccines, 4(1), 41. doi:10.1038/s41541-019-0134-4

Nimal, S., McCormick, A. L., Thomas, M. S., & Heath, A. W. (2005). An interferon gamma-gp120 fusion delivered as a DNA vaccine induces enhanced priming. Vaccine, 23(30), 3984–3990. doi:10.1016/j.vaccine.2005.01.160

O’Neill, L. A. J., & Bowie, A. G. (2010). Sensing and Signaling in Antiviral Innate Immunity. Curr Biol, 20(7), R328–R333. doi:https://doi.org/10.1016/j.cub.2010.01.044

Oda, S., Schroder, M., & Khan, A. R. (2009). Structural basis for targeting of human RNA helicase DDX3 by poxvirus protein K7. Structure, 17(11), 1528–1537. doi:10.1016/j.str.2009.09.005

Omura, N., Yoshikawa, T., Fujii, H., Shibamura, M., Inagaki, T., Kato, H., Egawa, K., Harada, S., Yamada, S., Takeyama, H., & Saijo, M. (2018). A Novel System for Constructing a Recombinant Highly-Attenuated Vaccinia Virus Strain (LC16m8) Expressing Foreign Genes and Its Application for the Generation of LC16m8-Based Vaccines against Herpes Simplex Virus 2. Jpn J Infect Dis, 71(3), 229–233. doi:10.7883/yoken.JJID.2017.458

Overton, E. T., Lawrence, S. J., Wagner, E., Nopora, K., Rosch, S., Young, P., Schmidt, D., Kreusel, C., De Carli, S., Meyer, T. P., Weidenthaler, H., Samy, N., & Chaplin, P. (2018). Immunogenicity and safety of three consecutive production lots of the non replicating smallpox vaccine MVA: A randomised, double blind, placebo controlled phase III trial. PLoS One, 13(4), e0195897. doi:10.1371/journal.pone.0195897

Pallett, M. A., Ren, H., Zhang, R. Y., Scutts, S. R., Gonzalez, L., Zhu, Z., Maluquer de Motes, C., & Smith, G. L. (2019). Vaccinia Virus BBK E3 Ligase Adaptor A55 Targets Importin-Dependent NF-kappaB Activation and Inhibits CD8(+) T-Cell Memory. J Virol, 93(10), e00051–00019. doi:10.1128/JVI.00051-19

Pantaleo, G., Janes, H., Karuna, S., Grant, S., Ouedraogo, G. L., Allen, M., Tomaras, G. D., Frahm, N., Montefiori, D. C., Ferrari, G., Ding, S., Lee, C., Robb, M. L., Esteban, M., Wagner, R., Bart, P. A., Rettby, N., McElrath, M. J., Gilbert, P. B., Kublin, J. G., Corey, L., & Network, N. H. V. T. (2019). Safety and immunogenicity of a multivalent HIV vaccine comprising envelope protein with either DNA or NYVAC vectors (HVTN 096): a phase 1b, double-blind, placebo-controlled trial. Lancet HIV, 6(11), e737–e749. doi:10.1016/S2352-3018(19)30262-0

Park, S., Choi, Y., Jung, N., Yu, Y., Ryu, K. H., Kim, H. S., Jo, I., Choi, B. O., & Jung, S. C. (2016). Myogenic differentiation potential of human tonsil-derived mesenchymal stem cells and their potential for use to promote skeletal muscle regeneration. Int J Mol Med, 37(5), 1209–1220. doi:10.3892/ijmm.2016.2536

Parrino, J., McCurdy, L. H., Larkin, B. D., Gordon, I. J., Rucker, S. E., Enama, M. E., Koup, R. A., Roederer, M., Bailer, R. T., Moodie, Z., Gu, L., Yan, L., Graham, B. S., & Team, V. R. C. S. (2007). Safety, immunogenicity and efficacy of modified vaccinia Ankara (MVA) against Dryvax challenge in vaccinia-naive and vaccinia-immune individuals. Vaccine, 25(8), 1513–1525. doi:10.1016/j.vaccine.2006.10.047

Paszkowski, P., Noyce, R. S., & Evans, D. H. (2016). Live-Cell Imaging of Vaccinia Virus Recombination. PLoS Pathog, 12(8), e1005824. doi:10.1371/journal.ppat.1005824

Perdiguero, B., & Esteban, M. (2009). The interferon system and vaccinia virus evasion mechanisms. J Interferon Cytokine Res, 29(9), 581–598. doi:10.1089/jir.2009.0073

Pham, A. M., Santa Maria, F. G., Lahiri, T., Friedman, E., Marie, I. J., & Levy, D. E. (2016). PKR Transduces MDA5-Dependent Signals for Type I IFN Induction. PLoS Pathog, 12(3), e1005489. doi:10.1371/journal.ppat.1005489

Pichlmair, A., Schulz, O., Tan, C. P., Rehwinkel, J., Kato, H., Takeuchi, O., Akira, S., Way, M., Schiavo, G., & Reis e Sousa, C. (2009). Activation of MDA5 requires higher-order RNA structures generated during virus infection. J Virol, 83(20), 10761–10769. doi:10.1128/JVI.00770-09

Pijlman, G. P., Suhrbier, A., & Khromykh, A. A. (2006). Kunjin virus replicons: an RNA-based, non-cytopathic viral vector system for protein production, vaccine and gene therapy applications. Expert Opin Biol Ther, 6(2), 135–145. doi:10.1517/14712598.6.2.135

Pittman, P. R., Hahn, M., Lee, H. S., Koca, C., Samy, N., Schmidt, D., Hornung, J., Weidenthaler, H., Heery, C. R., Meyer, T. P. H., Silbernagl, G., Maclennan, J., & Chaplin, P. (2019). Phase 3 Efficacy Trial of Modified Vaccinia Ankara as a Vaccine against Smallpox. N Engl J Med, 381(20), 1897–1908. doi:10.1056/NEJMoa1817307

Playfair, J. H., & De Souza, J. B. (1987). Recombinant gamma interferon is a potent adjuvant for a malaria vaccine in mice. Clin Exp Immunol, 67(1), 5–10.

Poo, Y. S., Nakaya, H., Gardner, J., Larcher, T., Schroder, W. A., Le, T. T., Major, L. D., & Suhrbier, A. (2014). CCR2 deficiency promotes exacerbated chronic erosive neutrophil-dominated chikungunya virus arthritis. J Virol, 88(12), 6862–6872. doi:10.1128/JVI.03364-13

Price, P. J., Luckow, B., Torres-Dominguez, L. E., Brandmuller, C., Zorn, J., Kirschning, C. J., Sutter, G., & Lehmann, M. H. (2014). Chemokine (C-C Motif) receptor 1 is required for efficient recruitment of neutrophils during respiratory infection with modified vaccinia virus Ankara. J Virol, 88(18), 10840–10850. doi:10.1128/JVI.01524-14

Prow, N. A., Hirata, T. D. C., Tang, B., Larcher, T., Mukhopadhyay, P., Alves, T. L., Le, T. T., Gardner, J., Poo, Y. S., Nakayama, E., Lutzky, V. P., Nakaya, H. I., & Suhrbier, A. (2019). Exacerbation of Chikungunya Virus Rheumatic Immunopathology by a High Fiber Diet and Butyrate. Front Immunol, 10, 2736. doi:10.3389/fimmu.2019.02736

Prow, N. A., Jimenez Martinez, R., Hayball, J. D., Howley, P. M., & Suhrbier, A. (2018a). Poxvirus-based vector systems and the potential for multi-valent and multi-pathogen vaccines. Expert Rev Vaccines, 17(10), 925–934. doi:10.1080/14760584.2018.1522255

Prow, N. A., Liu, L., McCarthy, M. K., Walters, K., Kalkeri, R., Geiger, J., Koide, F., Cooper, T. H., Eldi, P., Nakayama, E., Diener, K. R., Howley, P. M., Hayball, J. D., Morrison, T. E., & Suhrbier, A. (2020). The vaccinia virus based Sementis Copenhagen Vector vaccine against Zika and chikungunya is immunogenic in non-human primates. NPJ Vaccines, 5, 44. doi:10.1038/s41541-020-0191-8

Prow, N. A., Liu, L., Nakayama, E., Cooper, T. H., Yan, K., Eldi, P., Hazlewood, J. E., Tang, B., Le, T. T., Setoh, Y. X., Khromykh, A. A., Hobson-Peters, J., Diener, K. R., Howley, P. M., Hayball, J. D., & Suhrbier, A. (2018b). A vaccinia-based single vector construct multi-pathogen vaccine protects against both Zika and chikungunya viruses. Nat Commun, 9(1), 1230. doi:10.1038/s41467-018-03662-6

Quigley, M., Martinez, J., Huang, X., & Yang, Y. (2009). A critical role for direct TLR2-MyD88 signaling in CD8 T-cell clonal expansion and memory formation following vaccinia viral infection. Blood, 113(10), 2256–2264. doi:10.1182/blood-2008-03-148809

Rahman, M. T., Muppala, S., Wu, J., Krukovets, I., Solovjev, D., Verbovetskiy, D., Obiako, C., Plow, E. F., & Stenina-Adognravi, O. (2020). Effects of thrombospondin-4 on pro-inflammatory phenotype differentiation and apoptosis in macrophages. Cell Death Dis, 11(1), 53. doi:10.1038/s41419-020-2237-2

Reading, P. C., Symons, J. A., & Smith, G. L. (2003). A soluble chemokine-binding protein from vaccinia virus reduces virus virulence and the inflammatory response to infection. J Immunol, 170(3), 1435–1442. doi:10.4049/jimmunol.170.3.1435

Rehm, K. E., Connor, R. F., Jones, G. J., Yimbu, K., & Roper, R. L. (2010). Vaccinia virus A35R inhibits MHC class II antigen presentation. Virology, 397(1), 176–186. doi:10.1016/j.virol.2009.11.008

Reich, M., Liefeld, T., Gould, J., Lerner, J., Tamayo, P., & Mesirov, J. P. (2006). GenePattern 2.0. Nat Genet, 38(5), 500–501. doi:10.1038/ng0506-500

Rhyasen, G. W., & Starczynowski, D. T. (2015). IRAK signalling in cancer. Br J Cancer, 112(2), 232–237. doi:10.1038/bjc.2014.513

Robinson, J. T., Thorvaldsdóttir, H., Winckler, W., Guttman, M., Lander, E. S., Getz, G., & Mesirov, J. P. (2011). Integrative genomics viewer. Nat Biotechnol, 29(1), 24–26. doi:10.1038/nbt.1754

Rudd, P. A., Wilson, J., Gardner, J., Larcher, T., Babarit, C., Le, T. T., Anraku, I., Kumagai, Y., Loo, Y. M., Gale, M., Jr., Akira, S., Khromykh, A. A., & Suhrbier, A. (2012). Interferon response factors 3 and 7 protect against Chikungunya virus hemorrhagic fever and shock. J Virol, 86(18), 9888–9898. doi:10.1128/JVI.00956-12

Salem, S., Salem, D., & Gros, P. (2020). Role of IRF8 in immune cells functions, protection against infections, and susceptibility to inflammatory diseases. Hum Genet, 139(6-7), 707–721. doi:10.1007/s00439-020-02154-2

Samuelsson, C., Hausmann, J., Lauterbach, H., Schmidt, M., Akira, S., Wagner, H., Chaplin, P., Suter, M., O’Keeffe, M., & Hochrein, H. (2008). Survival of lethal poxvirus infection in mice depends on TLR9, and therapeutic vaccination provides protection. J Clin Invest, 118(5), 1776–1784. doi:10.1172/JCI33940

Sanada, T., Takaesu, G., Mashima, R., Yoshida, R., Kobayashi, T., & Yoshimura, A. (2008). FLN29 deficiency reveals its negative regulatory role in the Toll-like receptor (TLR) and retinoic acid-inducible gene I (RIG-I)-like helicase signaling pathway. J Biol Chem, 283(49), 33858–33864. doi:10.1074/jbc.M806923200

Sarkar, I., Garg, R., & van Drunen Littel-van den Hurk, S. (2019). Selection of adjuvants for vaccines targeting specific pathogens. Expert Rev Vaccines, 18(5), 505–521. doi:10.1080/14760584.2019.1604231

Schifanella, L., Barnett, S. W., Bissa, M., Galli, V., Doster, M. N., Vaccari, M., Tomaras, G. D., Shen, X., Phogat, S., Pal, R., Montefiori, D. C., LaBranche, C. C., Rao, M., Trinh, H. V., Washington-Parks, R., Liyanage, N. P. M., Brown, D. R., Liang, F., Lore, K., Venzon, D. J., Magnanelli, W., Metrinko, M., Kramer, J., Breed, M., Alter, G., Ruprecht, R. M., & Franchini, G. (2019). ALVAC-HIV B/C candidate HIV vaccine efficacy dependent on neutralization profile of challenge virus and adjuvant dose and type. PLoS Pathog, 15(12), e1008121. doi:10.1371/journal.ppat.1008121

Schrauf, S., Tschismarov, R., Tauber, E., & Ramsauer, K. (2020). Current Efforts in the Development of Vaccines for the Prevention of Zika and Chikungunya Virus Infections. Front Immunol, 11, 592. doi:10.3389/fimmu.2020.00592

Schroder, W. A., Le, T. T., Major, L., Street, S., Gardner, J., Lambley, E., Markey, K., MacDonald, K. P., Fish, R. J., Thomas, R., & Suhrbier, A. (2010). A physiological function of inflammation-associated SerpinB2 is regulation of adaptive immunity. J Immunol, 184(5), 2663–2670. doi:10.4049/jimmunol.0902187

Schwartz, L. M. (2008). Atrophy and programmed cell death of skeletal muscle. Cell Death Differ, 15(7), 1163–1169. doi:10.1038/cdd.2008.68

Schwartz, L. M. (2018). Skeletal Muscles Do Not Undergo Apoptosis During Either Atrophy or Programmed Cell Death-Revisiting the Myonuclear Domain Hypothesis. Front Physiol, 9, 1887. doi:10.3389/fphys.2018.01887

Scutts, S. R., Ember, S. W., Ren, H., Ye, C., Lovejoy, C. A., Mazzon, M., Veyer, D. L., Sumner, R. P., & Smith, G. L. (2018). DNA-PK Is Targeted by Multiple Vaccinia Virus Proteins to Inhibit DNA Sensing. Cell Rep, 25(7), 1953–1965 e1954. doi:10.1016/j.celrep.2018.10.034

Seubert, A., Monaci, E., Pizza, M., O’Hagan, D. T., & Wack, A. (2008). The adjuvants aluminum hydroxide and MF59 induce monocyte and granulocyte chemoattractants and enhance monocyte differentiation toward dendritic cells. J Immunol, 180(8), 5402–5412. doi:10.4049/jimmunol.180.8.5402

Shannon, P., Markiel, A., Ozier, O., Baliga, N. S., Wang, J. T., Ramage, D., Amin, N., Schwikowski, B., & Ideker, T. (2003). Cytoscape: a software environment for integrated models of biomolecular interaction networks. Genome Res, 13(11), 2498–2504. doi:10.1101/gr.1239303

Sharma, M., Krammer, F., Garcia-Sastre, A., & Tripathi, S. (2019). Moving from Empirical to Rational Vaccine Design in the ‘Omics’ Era. Vaccines (Basel), 7(3), 89. doi:10.3390/vaccines7030089

Simons, A. (2010). FastQC: A quality control tool for high throughput sequence data. Retrieved from https://www.bioinformatics.babraham.ac.uk/projects/fastqc

Smith, G. L., Benfield, C. T., Maluquer de Motes, C., Mazzon, M., Ember, S. W., Ferguson, B. J., & Sumner, R. P. (2013). Vaccinia virus immune evasion: mechanisms, virulence and immunogenicity. J Gen Virol, 94(Pt 11), 2367–2392. doi:10.1099/vir.0.055921-0

Smith, G. L., Talbot-Cooper, C., & Lu, Y. (2018). How Does Vaccinia Virus Interfere With Interferon? Adv Virus Res, 100, 355–378. doi:10.1016/bs.aivir.2018.01.003

So, T., & Ishii, N. (2019). The TNF-TNFR Family of Co-signal Molecules. Adv Exp Med Biol, 1189, 53–84. doi:10.1007/978-981-32-9717-3_3

Soehnlein, O., & Lindbom, L. (2010). Phagocyte partnership during the onset and resolution of inflammation. Nat Rev Immunol, 10(6), 427–439. doi:10.1038/nri2779

Stading, B., Ellison, J. A., Carson, W. C., Satheshkumar, P. S., Rocke, T. E., & Osorio, J. E. (2017). Protection of bats (Eptesicus fuscus) against rabies following topical or oronasal exposure to a recombinant raccoon poxvirus vaccine. PLoS Negl Trop Dis, 11(10), e0005958. doi:10.1371/journal.pntd.0005958

Stephen, J., Scales, H. E., Benson, R. A., Erben, D., Garside, P., & Brewer, J. M. (2017). Neutrophil swarming and extracellular trap formation play a significant role in Alum adjuvant activity. NPJ Vaccines, 2, 1. doi:10.1038/s41541-016-0001-5

Stuart, J. H., Sumner, R. P., Lu, Y., Snowden, J. S., & Smith, G. L. (2016). Vaccinia Virus Protein C6 Inhibits Type I IFN Signalling in the Nucleus and Binds to the Transactivation Domain of STAT2. PLOS Pathogens, 12(12), e1005955. doi:10.1371/journal.ppat.1005955

Subramanian, A., Tamayo, P., Mootha, V. K., Mukherjee, S., Ebert, B. L., Gillette, M. A., Paulovich, A., Pomeroy, S. L., Golub, T. R., Lander, E. S., & Mesirov, J. P. (2005). Gene set enrichment analysis: a knowledge-based approach for interpreting genome-wide expression profiles. Proc Natl Acad Sci U S A, 102(43), 15545–15550. doi:10.1073/pnas.0506580102

Suhrbier, A. (2019). Rheumatic manifestations of chikungunya: emerging concepts and interventions. Nat Rev Rheumatol, 15(10), 597–611. doi:10.1038/s41584-019-0276-9

Suschak, J. J., Wang, S., Fitzgerald, K. A., & Lu, S. (2016). A cGAS-Independent STING/IRF7 Pathway Mediates the Immunogenicity of DNA Vaccines. J Immunol, 196(1), 310–316. doi:10.4049/jimmunol.1501836

Sutherland, D. B., Ranasinghe, C., Regner, M., Phipps, S., Matthaei, K. I., Day, S. L., & Ramshaw, I. A. (2011). Evaluating vaccinia virus cytokine co-expression in TLR GKO mice. Immunol Cell Biol, 89(6), 706–715. doi:10.1038/icb.2010.157

Sutter, G. (2020). A vital gene for modified vaccinia virus Ankara replication in human cells. Proc Natl Acad Sci U S A, 117, 6289–6291.

Symons, J. A., Adams, E., Tscharke, D. C., Reading, P. C., Waldmann, H., & Smith, G. L. (2002). The vaccinia virus C12L protein inhibits mouse IL-18 and promotes virus virulence in the murine intranasal modelThe nucleotide sequence of the vaccinia virus strain Western Reserve C12L gene has been deposited at GenBank and assigned accession no. AF510447. J Gen Virol, 83(11), 2833–2844. doi:https://doi.org/10.1099/0022-1317-83-11-2833

Szklarczyk, D., Gable, A. L., Lyon, D., Junge, A., Wyder, S., Huerta-Cepas, J., Simonovic, M., Doncheva, N. T., Morris, J. H., Bork, P., Jensen, L. J., & Mering, C. V. (2019). STRING v11: protein-protein association networks with increased coverage, supporting functional discovery in genome-wide experimental datasets. Nucleic Acids Res, 47(D1), D607–D613. doi:10.1093/nar/gky1131

Szugye, H. S. (2020). Pediatric Rhabdomyolysis. Pediatr Rev, 41(6), 265–275. doi:10.1542/pir.2018-0300

Talbot, T. R., Peters, J., Yan, L., Wright, P. F., & Edwards, K. M. (2006). Optimal bandaging of smallpox vaccination sites to decrease the potential for secondary vaccinia transmission without impairing lesion healing. Infect Control Hosp Epidemiol, 27(11), 1184–1192. doi:10.1086/508827

Tameris, M., Geldenhuys, H., Luabeya, A. K., Smit, E., Hughes, J. E., Vermaak, S., Hanekom, W. A., Hatherill, M., Mahomed, H., McShane, H., & Scriba, T. J. (2014). The candidate TB vaccine, MVA85A, induces highly durable Th1 responses. PLoS One, 9(2), e87340. doi:10.1371/journal.pone.0087340

Tang, Q., Chakraborty, S., & Xu, G. (2018). Mechanism of vaccinia viral protein B14-mediated inhibition of IκB kinase β activation. J Biol Chem, 293(26), 10344–10352. doi:10.1074/jbc.RA118.002817

Tian, T., Dubin, K., Jin, Q., Qureshi, A., King, S. L., Liu, L., Jiang, X., Murphy, G. F., Kupper, T. S., & Fuhlbrigge, R. C. (2012). Disruption of TNF-alpha/TNFR1 function in resident skin cells impairs host immune response against cutaneous vaccinia virus infection. J Invest Dermatol, 132(5), 1425–1434. doi:10.1038/jid.2011.489

Tian, T., Jin, M. Q., Dubin, K., King, S. L., Hoetzenecker, W., Murphy, G. F., Chen, C. A., Kupper, T. S., & Fuhlbrigge, R. C. (2017). IL-1R Type 1-Deficient Mice Demonstrate an Impaired Host Immune Response against Cutaneous Vaccinia Virus Infection. J Immunol, 198(11), 4341–4351. doi:10.4049/jimmunol.1500106

Tolonen, N., Doglio, L., Schleich, S., & Krijnse Locker, J. (2001). Vaccinia virus DNA replication occurs in endoplasmic reticulum-enclosed cytoplasmic mini-nuclei. Mol Biol Cell, 12(7), 2031–2046. doi:10.1091/mbc.12.7.2031

Toor, I. S. (2019). Eosinophil deficiency promotes aberrant repair and adverse remodelling following acute myocardial infarction. bioRxiv. doi:https://doi.org/10.1101/750133

Torres-Domínguez, L. E., & McFadden, G. (2019). Poxvirus oncolytic virotherapy. Expert Opin Biol Ther, 19(6), 561–573. doi:10.1080/14712598.2019.1600669

Tscharke, D. C., & Suhrbier, A. (2005). From mice to humans - murine intelligence for human CD8+ T cell vaccine design. Expert Opin Biol Ther, 5(2), 263–271. doi:10.1517/14712598.5.2.263

Tsung, K., Yim, J. H., Marti, W., Buller, R. M., & Norton, J. A. (1996). Gene expression and cytopathic effect of vaccinia virus inactivated by psoralen and long-wave UV light. J Virol, 70(1), 165–171.

Unterholzner, L., Sumner, R. P., Baran, M., Ren, H., Mansur, D. S., Bourke, N. M., Randow, F., Smith, G. L., & Bowie, A. G. (2011). Vaccinia virus protein C6 is a virulence factor that binds TBK-1 adaptor proteins and inhibits activation of IRF3 and IRF7. PLoS Pathog, 7(9), e1002247. doi:10.1371/journal.ppat.1002247

van Slooten, M. L., Storm, G., Zoephel, A., Kupcu, Z., Boerman, O., Crommelin, D. J., Wagner, E., & Kircheis, R. (2000). Liposomes containing interferon-gamma as adjuvant in tumor cell vaccines. Pharm Res, 17(1), 42–48. doi:10.1023/a:1007514424253

Varikuti, S., Oghumu, S., Natarajan, G., Kimble, J., Sperling, R. H., Moretti, E., Kaplan, M. H., & Satoskar, A. R. (2016). STAT4 is required for the generation of Th1 and Th2, but not Th17 immune responses during monophosphoryl lipid A adjuvant activity. Int Immunol, 28(11), 565–570. doi:10.1093/intimm/dxw038

Waibler, Z., Anzaghe, M., Frenz, T., Schwantes, A., Pohlmann, C., Ludwig, H., Palomo-Otero, M., Alcami, A., Sutter, G., & Kalinke, U. (2009). Vaccinia virus-mediated inhibition of type I interferon responses is a multifactorial process involving the soluble type I interferon receptor B18 and intracellular components. J Virol, 83(4), 1563–1571. doi:10.1128/jvi.01617-08

Waibler, Z., Anzaghe, M., Ludwig, H., Akira, S., Weiss, S., Sutter, G., & Kalinke, U. (2007). Modified Vaccinia Virus Ankara Induces Toll-Like Receptor-Independent Type I Interferon Responses. J Virol, 81(22), 12102. doi:10.1128/JVI.01190-07

Walsh, G. M. (2020). Anti-IL-5 monoclonal antibodies for the treatment of asthma: an update. Expert Opin Biol Ther, 1–7. doi:10.1080/14712598.2020.1782381

Wang, J., Shah, D., Chen, X., Anderson, R. R., & Wu, M. X. (2014). A micro-sterile inflammation array as an adjuvant for influenza vaccines. Nat Commun, 5, 4447. doi:10.1038/ncomms5447

Wang, L., Liebmen, M. N., & Wang, X. (2017). Roles of Mitochondrial DNA Signaling in Immune Responses. Adv Exp Med Biol, 1038, 39–53. doi:10.1007/978-981-10-6674-0_4

Wang, L., Wang, S., & Li, W. (2012). RSeQC: quality control of RNA-seq experiments. Bioinformatics, 28(16), 2184–2185. doi:10.1093/bioinformatics/bts356

Weller, P. F., & Spencer, L. A. (2017). Functions of tissue-resident eosinophils. Nat Rev Immunol, 17(12), 746–760. doi:10.1038/nri.2017.95

Wilson, J. A., Prow, N. A., Schroder, W. A., Ellis, J. J., Cumming, H. E., Gearing, L. J., Poo, Y. S., Taylor, A., Hertzog, P. J., Di Giallonardo, F., Hueston, L., Le Grand, R., Tang, B., Le, T. T., Gardner, J., Mahalingam, S., Roques, P., Bird, P. I., & Suhrbier, A. (2017). RNA-Seq analysis of chikungunya virus infection and identification of granzyme A as a major promoter of arthritic inflammation. PLoS Pathog, 13(2), e1006155. doi:10.1371/journal.ppat.1006155

Wolferstatter, M., Schweneker, M., Spath, M., Lukassen, S., Klingenberg, M., Brinkmann, K., Wielert, U., Lauterbach, H., Hochrein, H., Chaplin, P., Suter, M., & Hausmann, J. (2014). Recombinant modified vaccinia virus Ankara generating excess early double-stranded RNA transiently activates protein kinase R and triggers enhanced innate immune responses. J Virol, 88(24), 14396–14411. doi:10.1128/JVI.02082-14

Wyatt, L. S., Xiao, W., Americo, J. L., Earl, P. L., & Moss, B. (2017). Novel Nonreplicating Vaccinia Virus Vector Enhances Expression of Heterologous Genes and Suppresses Synthesis of Endogenous Viral Proteins. mBio, 8(3). doi:10.1128/mBio.00790-17

Yang, C., Mai, H., Peng, J., Zhou, B., Hou, J., & Jiang, D. (2020). STAT4: an immunoregulator contributing to diverse human diseases. Int J Biol Sci, 16(9), 1575–1585. doi:10.7150/ijbs.41852

Yang, M. Q., Du, Q., Varley, P. R., Goswami, J., Liang, Z., Wang, R., Li, H., Stolz, D. B., & Geller, D. A. (2018). Interferon regulatory factor 1 priming of tumour-derived exosomes enhances the antitumour immune response. Br J Cancer, 118(1), 62–71. doi:10.1038/bjc.2017.389

Yokota, S.-I., Okabayashi, T., & Fujii, N. (2010). The battle between virus and host: modulation of Toll-like receptor signaling pathways by virus infection. Mediators of inflammation, 2010, 184328. doi:10.1155/2010/184328

Yoshida, H., & Hunter, C. A. (2015). The immunobiology of interleukin-27. Annu Rev Immunol, 33, 417–443. doi:10.1146/annurev-immunol-032414-112134

Yoshimoto, Y., Ikemoto-Uezumi, M., Hitachi, K., Fukada, S. I., & Uezumi, A. (2020). Methods for Accurate Assessment of Myofiber Maturity During Skeletal Muscle Regeneration. Front Cell Dev Biol, 8, 267. doi:10.3389/fcell.2020.00267

Zhang, H., Liu, Z., & Liu, S. (2016). HMGB1 induced inflammatory effect is blocked by CRISPLD2 via MiR155 in hepatic fibrogenesis. Mol Immunol, 69, 1–6. doi:10.1016/j.molimm.2015.10.018

Zhao, C., & Zhao, W. (2019). TANK-binding kinase 1 as a novel therapeutic target for viral diseases. Expert Opin Ther Targets, 23(5), 437–446. doi:10.1080/14728222.2019.1601702

